# Ginsenoside Ro ameliorates diabetic cardiomyopathy by maintaining ciliary homeostasis and enhancing antioxidation

**DOI:** 10.64898/2026.06.22.733885

**Authors:** Zhuangzhuang Yang, Yi Guo, Boyu Guan, Xiying Guo, Yafang Shang, Yongsong Tang, Chenchen Zhao, Peng Wang, Zhanhong Ren

## Abstract

**Objective:** To investigate and clarify the role of Ginsenoside Ro (GRo) in diabetic cardiomyopathy (DiaCM) and to elucidate the molecular mechanism by which GRo ameliorates DiaCM.

**Methods:** ① **The construct of type 2 diabetic mouse model**. The bought C57BL/6 male mice were housed in a specific pathogen-free (SPF) animal facility and randomly divided into control, STZ (model), STZ + GRo, and control+GRo groups. The STZ (model) and STZ + GRo groups were fed a high-fat and high-glucose diet combined with intraperitoneal injection of streptozotocin (STZ). The control and control + GRo groups were fed a normal diet, while the control + GRo and STZ + GRo groups were treated with GRo via oral gavage. Then, all groups were evaluated for cardiac function and structure by small animal echocardiography and histological staining including hematoxylin and eosin (HE) and Masson’s trichrome staining to screen and confirm diabetic cardiomyopathy in mice. Finally, immunofluorescence staining of cilia in mouse heart tissue was performed to determine whether GRo inhibits abnormal ciliary growth. ② **The construct of cell models**. First, the CCK-8 (Cell Counting Kit-8) assay was used to separately evaluate the cytotoxicity of GRo and the combination of TGF-β1 and PA in myocardial fibroblasts and cardiomyocytes. Subsequently, mouse myocardial fibroblast lines (MCFs) were treated with transforming growth factor-beta 1 (TGF-β1), and H9c2 cardiomyocytes were treated with palmitic acid (PA). Both cell types then received the GRo treatment. ③ **Molecular and cellular testing**. Firstly, we measured serum levels of cardiac injury markers (CK-MB, MYO, and TNNI3), glutathione (GSH), and malondialdehyde (MDA). Secondly, we examined the expression of myocardial fibrosis-related genes (*Col1a1*, etc.), myocardial hypertrophy markers (*Nppa*, etc.), cilia-specific genes (*Pkd1*, etc.), and oxidative stress-related genes (*Nrf2*, etc.) in both animal and cell samples by Western blotting and RT-qPCR. Finally, we used immunofluorescence staining of myocardial fibroblasts to detect cilia length and phalloidin staining of cardiomyocytes to measure their cross-sectional area. ④ **The correlation mechanism**. Firstly, the cilia-specific inhibitory drug HIP-4 was used to disrupt cilia homeostasis by inhibiting cilia growth. Secondly, small activating RNA (saRNA) was used to upregulate the *Pkd1* gene to verify whether GRo exerts its anti-fibrotic effects through the inhibition of PC1.

**Results:** ① **Animal level**. A diabetic cardiomyopathy mouse model was successfully established by combining STZ injection with a high-fat and high-glucose diet, and treatment with GRo significantly ameliorated the associated symptoms. ② **Cellular level.** We successfully established a myocardial fibrosis model by treating myocardial fibroblasts with TGF-β1, and a myocardial hypertrophy model by treating cardiomyocytes with PA. Immunofluorescence staining demonstrated that GRo significantly decreased cilia length in the fibrosis model, while phalloidin staining showed that GRo significantly attenuated the increase in cardiomyocyte cross-sectional area. ③ **Molecular level**. Compared with the model group, GRo treatment significantly reduced serum levels of cardiac injury markers (CK-MB, MYO and TNNI3), glutathione (GSH) and malondialdehyde (MDA). Western blotting and RT-qPCR analyses of both animal and cell samples revealed that GRo markedly alleviated indicators of myocardial fibrosis and hypertrophy, while also suppressing cilia-specific genes and oxidative stress-related genes. Overall, GRo significantly ameliorated the markers associated with myocardial fibrosis and hypertrophy, and inhibited cilia-specific protein expression as well as oxidative stress parameters. ④ **The correlation mechanism**. The cilia-specific drug hedgehog pathway inhibitor 4 (HPI-4) was used to revealed that cilia homeostasis is closely linked to myocardial fibrosis and shortened cilia inhibit the fibrosis progression. Furthermore, upregulation of the *Pkd1* gene by small activating RNA demonstrated that PC1 overexpression abrogates the therapeutic effect of GRo. Finally, GRo can alleviate DiaCM.

## Introduction

Diabetes has emerged as one of the fastest-growing global health crises in the 21st century. In 2024, an estimated 589 million adults aged 20–79 years were living with diabetes worldwide, representing a prevalence of 11.1%, and it is projected to rise to 853 million by 2050^[1–5]^. Chronic hyperglycemia can lead to a spectrum of multisystem complications including diabetic cardiomyopathy (DiaCM), nephropathy, retinopathy, neuropathy and foot disease**^Error! Reference source not found.^**. Notably, DiaCM represents the leading cause of death in individuals with diabetes, which accounts for approximately 70% of diabetes-related mortality^[12–13]^. It is characterized as structural and functional abnormalities of the heart induced by hyperglycemia, independent of hypertension, coronary artery disease and other cardiac pathologies^[14–16]^. Clinically, it often manifests initially as diastolic dysfunction, and progresses to cardiac enlargement and impaired systolic function, which ultimately leads to heart failure^[17]^. The central pathological feature of DiaCM is myocardial remodelling which is a key driver of heart failure^[18]^. Therefore, inhibiting myocardial remodelling is beneficial to ameliorate DiaCM and prevent heart failure.

The two main events of myocardial remodeling include myocardial fibrosis and hypertrophy^[19–22]^. It has been extensively and thoroughly studied that the persistent activation of the renin-angiotensin-aldosterone system (RAAS), overactivation of the transforming growth factor-β (TGF-β) signaling pathway and infiltration of inflammatory factors can result in myocardial fibrosis^[23–35]^. Recent studies have shown that abnormal ciliary homeostasis also can induce myocardial fibrosis^[36–40]^. Cilia on the surface of myocardial fibroblasts serve as important sensory receptors and their abnormal growth can lead to myocardial fibrosis^[41–42]^. It is reported that polycystin-1 (PC1), a type of cilia-specific membrane signalling proteins encoded by polycystic kidney disease 1 (*Pkd1*), can upregulate the TGF-β/SMAD signalling pathway and drive the expression of fibrosis-related genes such as *Col1a1*, *Col1a3* and *Acta2*, which ultimately accelerates extracellular matrix deposition^[43–47]^. In addition, hyperglycaemia-induced oxidative stress can serve as the primary driver of myocardial hypertrophy^[48–51]^. In the heart of diabetic mouse, markedly elevated levels of reactive oxygen species (ROS) can suppress the Nrf2/SLC7A11/GPX4 pathway, which thereby incudes the lipid peroxidation and myocardial hypertrophy^[52–60]^.

Current therapeutic drugs such as sodium–glucose cotransporter 2 (SGLT2) inhibitors, glucagon-like peptide-1 (GLP-1) receptor agonists and β-blockers for DiaCM primarily focus on the glycaemic control and improvement of cardiac function^[61–77]^. However, their long-term use is often associated with adverse effects such as hypoglycaemia, hypotension and renal impairment, and their efficacy in reversing myocardial remodelling remains limited^[78–80]^. Compared with them, ginsenosides have high safety, strong metabolic adaptability, a multi-target, multi-pathway holistic regulatory mechanism, and excellent anti-inflammatory and antioxidant capacity^[81–83]^. Ginsenosides, a type of natural bioactive constituents derived from *Panax* species, are classified into oleanane-type and dammarane-type and they exert pronounced cardiovascular protective effects^[84–86]^. Compared with the dammarane type, the oleanane type including GRo has been less studied in Cardiovascular disease^[87-90]^. Previous studies have shown that GRo possesses distinct pharmacokinetic advantages including favourable water solubility, low first-pass effect, high bioavailability, Strong anti-inflammatory, antioxidant, neuroprotective, vascular protective and antibacterial properties^[91–93]^. Here, our results have shown that GRo can maintain ciliary homeostasis by downregulating cilia-specific proteins such as PC1, which thereby inhibits myocardial fibrosis. In addition, it can suppress oxidative stress by upregulating the Nrf2–SLC7A11/GPX4 antioxidant pathway, which thereby suppresses myocardial hypertrophy. Therefore, it can ameliorate DiaCM by blocking myocardial remodeling. Our study not only contributes to furtherly investigate the molecular mechanisms underlying the occurrence and development of DiaCM, but also provides novel potential therapeutic strategies for the prevention and treatment of DiaCM.

## Materials and methods

### Experimental animals

Specific-pathogen-free (SPF) male C57BL/6 mice (36 animals, 8 weeks of age, body weight 20–22 g) were obtained from Hunan Slyke Jingda Laboratory Animal Co., Ltd. (animal license no. SCXK (Xiang) 2021-0002). To induce diabetes, mice were given streptozotocin (STZ, 50 mg/kg) by gavage once daily for five consecutive days and were maintained on a high-sugar, high-fat diet. After the onset of diabetes was confirmed, the animals received GRo (10 mg/kg, intraperitoneal injection) daily for two months. All mice were housed at Hubei University of Science and Technology under conditions simulating the normal living environment: the room was kept well ventilated and hygienic, with a controlled temperature of 23 ± 3 °C, relative humidity of 55–64%, a 12-h light/dark cycle, and ad libitum access to food and drinking water.

### Establishment of animal models

Thirty-six eight-week-old male C57BL/6J mice (body weight 22–24 g) were purchased and allowed to acclimatize for one week. Mice were then randomly divided into a normal control group (n = 18) and a type 2 diabetic model group (n = 18). STZ was dissolved in 0.1 mol/L sodium citrate buffer (pH 4.5) and administered intraperitoneally to the model group at a dose of 50 mg/kg under light-protected conditions once daily for five consecutive days. One week after the final injection, fasting blood glucose was measured; a fasting blood glucose level exceeding 11.1 mmol/L was considered indicative of successful induction of diabetes. Following confirmation of diabetes, all mice were maintained on a high-sugar, high-fat diet for one month, and at week 7, fasting blood glucose was reassessed to confirm that levels remained above 11.1 mmol/L in the type 2 diabetic mice. The normal control mice were then randomly subdivided into a control group and a control + GRo group, while the type 2 diabetic mice were randomly subdivided into a STZ group and a STZ + GRo group. Starting at week 8, GRo (10 mg/kg) was administered by oral gavage once daily for 2 months. At week 16, treatment was discontinued and cardiac function was evaluated by echocardiography; mice were sacrificed and tissues were collected at week 18.

### Determination of cardiac function

Cardiac function was assessed after treatment using a high-resolution small-animal ultrasound imaging system. On the night before the examination, the hair on the left chest of each mouse was carefully removed to ensure close contact between the ultrasound transducer and the skin and to obtain clearer images. Mice were anesthetized with isoflurane, placed in the supine position on the platform, and their limbs were gently secured. The transducer was positioned on the left hemithorax, two-dimensional and M-mode images were acquired, and the following parameters were measured offline: left ventricular ejection fraction (LVEF%), left ventricular fractional shortening (LVFS%), left ventricular posterior wall thickness at end-diastole (LVPW; d), left ventricular posterior wall thickness at end-systole (LVPW; s), left ventricular anterior wall thickness at end-diastole (LVAW; d) and left ventricular anterior wall thickness at end-systole (LVAW; s)

### Hematoxylin and eosin (H&E) staining and Masson’s trichrome staining

Hearts were fixed for 12 h and subsequently sent to Servicebio (Wuhan, China) for paraffin embedding and hematoxylin and eosin (H&E) staining. Stained sections were imaged by panoramic scanning using a Zeiss microscope equipped with a 1× objective. To evaluate the extent of myocardial fibrosis, paraffin-embedded cardiac sections were stained with Masson’s trichrome and imaged under a Zeiss microscope at 40× magnification.

### Tissue section staining

Paraffin-embedded mouse heart sections were routinely deparaffinized and rehydrated. Antigen retrieval was performed by high-pressure and high-temperature treatment in EDTA buffer (pH=9.0, Hubei BioAus Biotech Co., Ltd.) for 1.5 min. After natural cooling, the sections were washed with TBST. Endogenous peroxidase activity was blocked with 3% H₂O₂ (Sinopharm Chemical Reagent Co., Ltd.) at room temperature for 30 min, followed by blocking with 10% normal goat serum (Wuhan Boster Biological Technology Co., Ltd.) at 37°C for 30 min. The primary antibody, mouse anti-ARL13B (1:2000, Abcam), was applied overnight at 4°C. The next day, the sections were incubated with HRP-conjugated goat anti-mouse secondary antibody (1:2000, Abcam) at 37°C for 45 min, then with iFluor® 488 tyramide (TSA, 1:600 in 0.003% H₂O₂) at room temperature for 10 min, followed by TBST washes. Antibody stripping was carried out by microwave heating (medium power) in citrate buffer (pH=6.0) for 5 min, after cooling, the sections were washed again with TBST. Nuclei were counterstained with DAPI (1:500) for 5 min. Finally, the sections were mounted and images were acquired using a fluorescence microscope.

### Cell culture and viability assays

The cell culture methods were based on previous literature and H9c2 and MCFS cells were cultured in Dulbecco’s Modified Eagle Medium (DMEM) medium for subculturing and plating. The cytotoxicity of GRo against H9c2 and MCFS cells was evaluated using the CCK-8 (Wuhan Servicebio) assay. Through co-administration of PA + GRo and TGF-β1 + GRo, it was ultimately determined that GRo exhibited the lowest cytotoxicity at a concentration of 40 μM (**Supplementary Fig. 1A–B, Supplementary Fig. 4A–E**).

### The establishment of cell models

Mouse myocardial fibroblast lines (MCFs) (purchased from Wuhan Procell Life Science & Technology Co., Ltd.) were used to establish the cell model of myocardial fibrosis induced by 10 ng/mL transforming growth factor-β1 (TGF-β1, TargetMol). MCFS were divided into four groups: control, TGF-β1, TGF-β1 + GRo and control + GRo. H9c2 cells (purchased from Wuhan Procell Life Science & Technology Co., Ltd.) were assigned to the same four groups and treated by 200 μM palmitic acid (PA, Yuanye Bio) to induce cardiomyocyte hypertrophy.

### RNA extraction and real-time PCR

Total RNA was extracted from cells and tissues using the TRIzol method (Thermo Fisher Scientific). The mRNA expression levels of the myocardial hypertrophy-related markers, fibrosis-related markers, cilia-specific genes, and oxidative stress (**Supplementary Table 1**) in myocardial tissue and cardiomyocytes were quantified by real-time quantitative PCR, with *GAPDH* serving as the internal control. All primers were synthesized by Sangon Biotech (Wuhan, China). The amplification protocol was as follows: initial denaturation at 95 °C for 30 s, followed by 30 cycles of denaturation at 95 °C for 10 s and annealing/extension at 60 °C for 30 s. Melting curve analysis was performed by increasing the temperature from 65 °C to 95 °C (0.5 °C increment per cycle). Relative gene expression was calculated using the 2^^(−ΔΔCq)^ method.

### Cells were transfected with small activating RNA (saRNA)

In a 1.5 mL microcentrifuge tube, 200 μL of the DMEM (Wuhan Servicebio) was mixed with an appropriate volume of GP-transfect-Mate transfection reagent (SignaGen). In a separate 1.5 mL tube, 200 μL of DMEM was combined with an appropriate amount of plasmid carrying the *Pkd1* gene fragment (**Supplementary Table 2**). Both tubes were left to stand in a biosafety cabinet for 5 min to allow the contents to mix thoroughly. The diluted transfection reagent was then added to the diluted plasmid solution, gently mixed by pipetting, and incubated for a further 15 min at room temperature in the biosafety cabinet. During this incubation, the DMEM was removed from the 6-well plates containing seeded cells, the cells were washed twice with PBS, and 1.6 mL of fresh DMEM was added to each well. After incubation, 400 μL of the transfection complex mixture was added dropwise to the corresponding wells, and the plate was gently swirled in a crosswise motion to ensure even distribution. Cells were incubated at 37 °C with 5% CO₂ for 6–8 h, after which the DMEM containing the transfection complexes was replaced with 2 mL of the serum-containing DMEM containing. Cells were further cultured for 24 h before subsequent experiments.

### Western blotting

Total protein was extracted from cardiac tissues and cells using RIPA lysis buffer (Thermo Fisher Scientific). Equal amounts of protein (30 μg) were separated by sodium dodecyl sulfate–polyacrylamide gel electrophoresis (SDS–PAGE, Bio-Rad) and transferred onto polyvinylidene difluoride (PVDF) membranes. Membranes were blocked with 5% non-fat dry milk and subsequently incubated with primary antibodies overnight at 4 °C, followed by incubation with the corresponding secondary antibodies. The following primary antibodies were used: COL I, COL III, α-SMA, SMAD2, SMAD3, phospho-SMAD2 (P-SMAD2), and phospho-SMAD3 (P-SMAD3) antibodies (Wuhan ABclonal Biotechnology Co., Ltd.); PC1 and KIF3A antibodies (Zen BioScience); PC2 antibody (Wuhan Sanying/Proteintech); IFT88 antibody (Zen BioScience); and Nrf2, GPX4, P53, and SLC7A11 antibodies (Zen BioScience). Primary antibodies were diluted in primary antibody dilution buffer (Biosharp).

### Immunofluorescence staining

When cells reached 50–60% confluence, they were serum-starved in DMEM containing 1% fetal bovine serum (FBS) for 12 h, after which the designated treatments (model induction and drug administration) were applied as described above. Twenty-four hours later, cells were washed three times with PBS and fixed with 1% paraformaldehyde for 20 min at room temperature. After three additional washes with PBS (5 min each), cells were permeabilized with 0.1% Triton X-100 in PBS for 5 min and washed again three times with PBS. Cells were then blocked with 1% goat serum for 1 h at room temperature. Following blocking, cells were washed three times with PBS containing 0.1% Triton X-100 (5 min per wash) and incubated with an anti-acetylated-α-tubulin primary antibody (1:100 dilution, 70 μL per sample, Santa cruz biotechnology, INC) for 1.5 h at room temperature in the dark. After incubation, cells were washed three times with PBS containing 0.1% Triton X-100. The slides were mounted with DAPI-containing mounting medium (100 μL) for 10 min in the dark, sealed with parafilm, and stored protected from light. Images were acquired using a confocal microscope to visualize cilia morphology.

### Molecular docking

The structure of the target compound was drawn using ChemDraw. The small-molecule ligand (GRo) was processed with ChemDraw 3D to generate its three-dimensional conformation, which was then converted into the PDBQT format for molecular docking. The protein structure of the PC1 receptor was retrieved from the RCSB Protein Data Bank (PDB). Using PyMOL (version 2.3.4), water molecules and native ligands were removed from the receptor structure. AutoDock Tools was subsequently employed to add hydrogen atoms and assign Gasteiger charges to the receptor. Both the PC1 receptor and GRo were ultimately converted into the PDBQT format. Molecular docking between the PC1 receptor and GRo was performed using AutoDock Vina (version 1.1.2). The docking results were analyzed with PLIP (Protein–Ligand Interaction Profiler) and visualized using PyMOL.

### Statistical analysis

ImageJ (NIH) and GraphPad Prism 9 (GraphPad Software) were used for image processing and statistical analysis. Selected graphical elements were created using BioGDP. All quantitative data are presented as mean ± SEM. Differences between two groups were assessed using the two-tailed unpaired Student’s t-test, while comparisons among three or more groups were performed by one-way analysis of variance (one-way ANOVA). A *P* value of less than 0.05 was considered statistically significant.

## Results

### GRo improves cardiac structure and function in type 2 diabetic mice

To evaluate the therapeutic effect of GRo in diabetic cardiomyopathy (DiaCM), the type 2 diabetic mice model was established by using the STZ injection combined with a high-sugar and high-fat diet, and the mice were treated with GRo by oral gavage for 8 weeks. Compared with the control group, fasting blood glucose levels in the STZ group were persistently elevated and eventually stabilized (**Fig. 1A**). Fasting blood glucose was significantly lower in the STZ + GRo group than that in the STZ group, which indicates that the GRo treatment can attenuates STZ-induced sustained fasting hyperglycemia (**Fig. 1A**). The STZ group displayed marked cardiac enlargement relative to the control group (**Fig. 1B**). Notably, the STZ + GRo group exhibited a significantly reduced heart size compared with the STZ group (**Fig. 1B**). Echocardiographic analysis further demonstrated that STZ-treated mice developed myocardial structural abnormalities and dysfunction. (**Fig. 1C**). Compared with the STZ group, GRo can effectively decrease the heart-weight–to–body-weight ratio (***p*=0.0047**), left ventricular posterior wall thickness at end-diastole (LVPW; d) (***p*=0.0040**), left ventricular posterior wall thickness at end-systole (LVPW; s) (***p*=0.0003**), left ventricular anterior wall thickness at end-diastole (LVAW; d) (***p*=0.0002**), left ventricular anterior wall thickness at end-systole (LVAW; s) (***p*=0.0016**) and restore the ejection fraction (EF) (***p*=0.0055**), fractional shortening (FS) (***p*=0.0071**) (**Fig. 1C–J**). It is indicated that GRo exerts cardioprotective effects through controling the hyperglycemia and reversing myocardial morphological abnormalities and dysfunction in DiaCM.

**Fig. 1.**
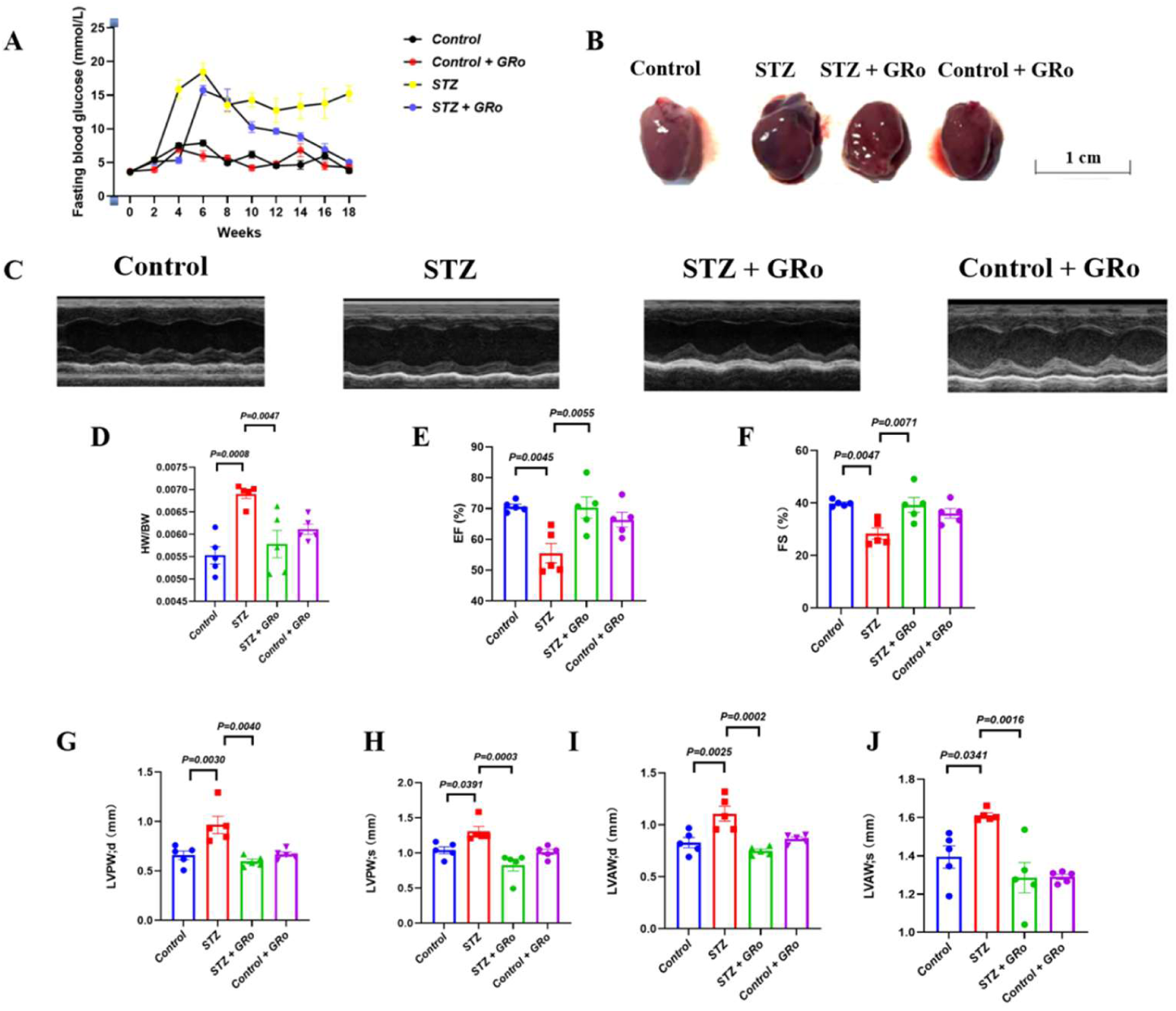
GRo improves cardiac structure and function in type 2 diabetic mice. **A**: Curves of fasting blood glucose changes in control, STZ, STZ + GRo and control + GRo groups over 18 weeks. **B**: Whole heart images of control, STZ, STZ + GRo and control + GRo groups. Scale bar, 1 cm. **C**: Representative left ventricular M-mode echocardiographic images. **D-J**: Quantitative indicators of cardiac function and structure from control, STZ, STZ + GRo and control + GRo groups: heart weight/body weight ratio (HW/BW) (**D**), ejection fraction (EF) (**E**), fractional shortening (FS) (**F**), left ventricular posterior wall thickness at end-diastole (LVPW; d) (**G**), left ventricular posterior wall thickness at end-systole (LVPW; s) (**H**), left ventricular anterior wall thickness at end-diastole (LVAW; d) (**I**) and left ventricular anterior wall thickness at end-systole (LVAW; s) (**J**). Data are presented as mean±SEM, n=5.

### GRo attenuates myocardial remodeling and reduces cardiac injury in type 2 diabetic mice

To further clarify the protective effects of GRo in DiaCM, myocardial structure and interstitial fibrosis were assessed by using hematoxylin and eosin (H&E) and Masson’s trichrome staining methods. Compared with the control group, the STZ group displayed marked cardiomyocyte hypertrophy and myocardial fibrosis (**Fig. 2A**). Remarkably, the GRo treatment in the STZ + GRo group reversed the STZ-induced cardiomyocyte hypertrophy and myocardial fibrosis (**Fig. 2A**). Quantitative analysis of Masson’s trichrome-stained sections revealed that the extent of myocardial fibrosis was substantially greater in the STZ group relative to the control group (***P* < 0.0001**) while the GRo administration significantly reduced the collagen deposition area compared with the STZ group (***P* =0.0007) (Fig. 2B**). Serological analyses demonstrated that the STZ group exhibited markedly elevated levels of three cardiac injury markers (CK-MB, TNNI3 and MYO) (***P* < 0.001**) and malondialdehyde (MDA) (***P* =0.0188**), along with a pronounced decrease in glutathione (GSH) level (***P* < 0.0001**) compared with the control group (**Fig. 2C–G**). Notably, the GRo treatment in the STZ + GRo group downregulated three cardiac injury markers compared with the STZ group (***P* < 0.01, Fig. 2C–E**). In addition, GRo significantly restored myocardial redox homeostasis (***P* < 0.05, Fig. 2F-G**). It is demonstrated that GRo exerts anti-myocardial remodeling, anti-cardiac injury and anti-oxidative stress in DiaCM.

**Fig. 2.**
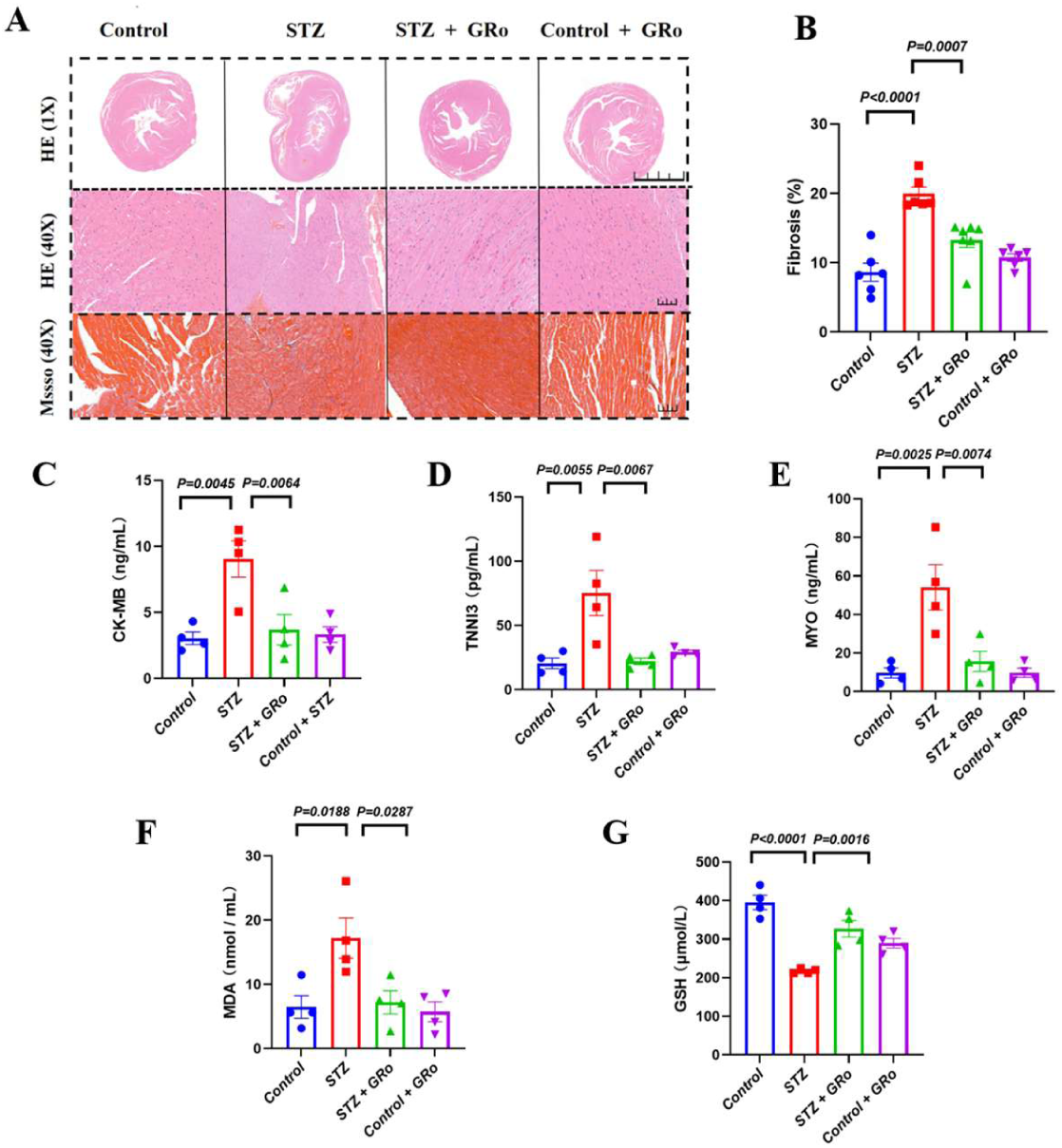
GRo attenuates myocardial remodeling and reduces cardiac injury in type 2 diabetic mice. **A**: Representative images of H&E and Masson’s trichrome staining from control, STZ, STZ + GRo and control + GRo groups. Scale bars, 1 000 μm and 20 μm for H&E, 20 μm for Masson’s trichrome staining. **B**: Quantification of cardiac fibrosis area in Masson’s trichrome staining. Data are presented as mean±SEM, n=6. **C-E**: Quantification of cardiac injury markers in serum from control, STZ, STZ + GRo and control + GRo groups: creatine kinase isoenzyme (CK-MB) (**C**), cardiac troponin I (TNNI3) (**D**), and myoglobin (MYO) (**E**). Data are presented as mean±SEM, n=4. **F-G**: Quantification of oxidative stress indicators in myocardial tissues from control, STZ, STZ + GRo and control + GRo groups: malondialdehyde (MDA) (**F**) and glutathione (GSH) (**G**). Data are presented as mean±SEM, n=4.

### GRo alleviates myocardial fibrosis in type 2 diabetic mice by downregulating the TGF-β/SMAD signaling pathway

To reveal the roles of GRo in myocardial fibrosis, we examined the levels of fibrosis-related markers (COLI, COLIII and α-SMA) and the activation status of the TGF-β/SMAD signalling in cardiac tissues. The results of western blot and RT-qPCR assays showed that the protein and mRNA levels of fibrosis-related markers were significantly upregulated in the STZ group compared with the control group (***P* < 0.05, Fig. 3A–I**). Compared with the STZ group, the GRo treatment in the STZ + GRo group markedly downregulated the fibrosis-related markers (***P* < 0.05, Fig. 3A–F**). Furtherly, the results of western blot assays showed that the TGF-β1 and phosphorylation of SMAD2/3 were significantly elevated in the STZ group compared with the control group (***P* < 0.05, Fig. 3J–O)**, while the GRo administration in the STZ + GRo group significantly downregulated the TGF-β1 and inhibited the phosphorylation of SMAD2/3 compared with the STZ group (***P* < 0.05, Fig. 3J–O**). It is demonstrated that GRo can attenuate myocardial fibrosis by reducing the excessive collagen deposition and downregulating the TGF-β/SMAD signalling in DiaCM.

**Fig 3.**
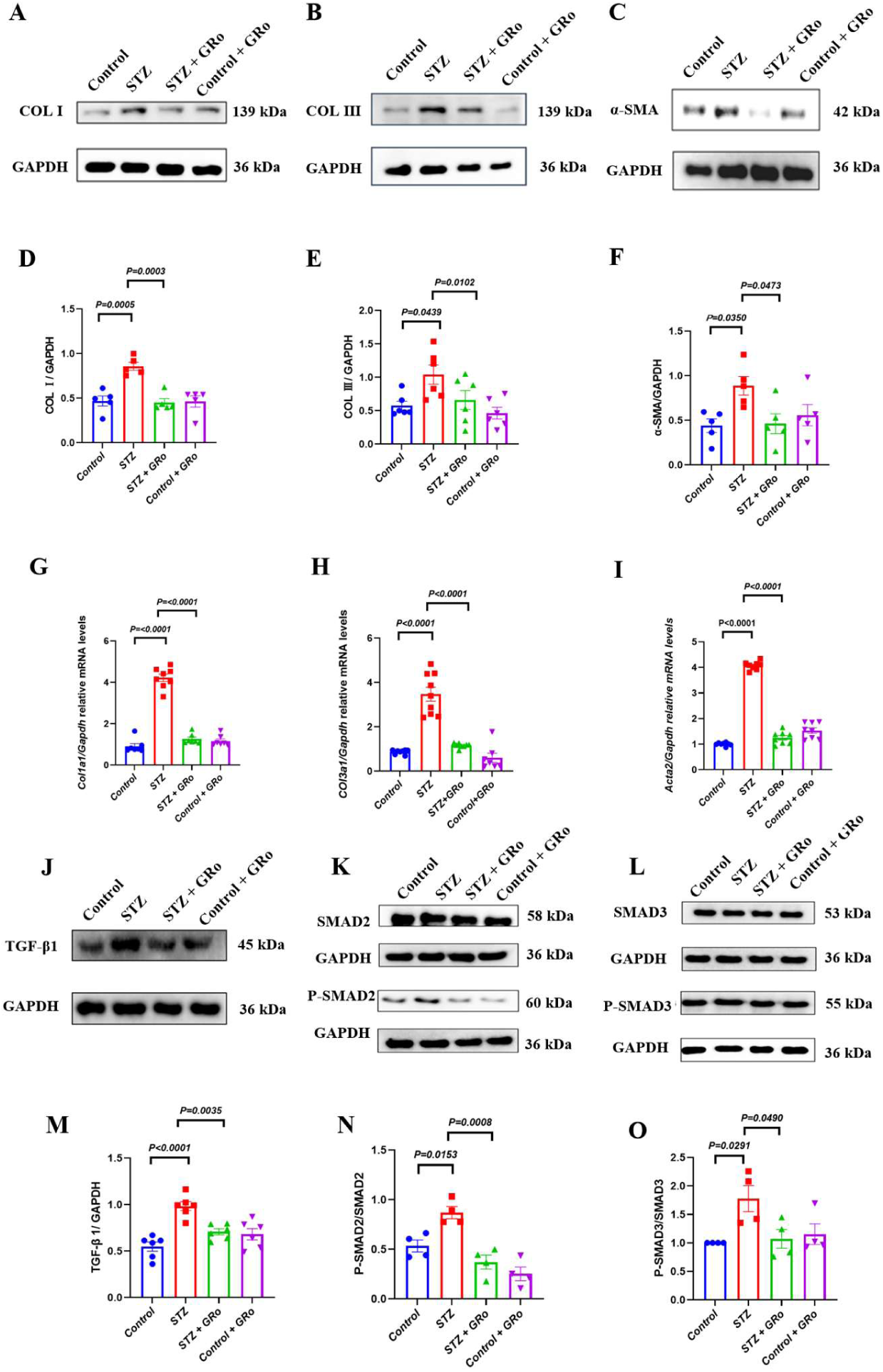
GRo alleviates myocardial fibrosis in type 2 diabetic mice by downregulating the TGF-β/SMAD signaling pathway. **A-C**: Representative western blot images of COL I, COL III and α-SMA in cardiac tissues from control, STZ, STZ + GRo and control + GRo groups. **D**: Quantification of western blots of COL I. n=5. **E**: Quantification of western blots of COL III. n=6. **F**: Quantification of western blots of α-SMA. n=5. **G-I**: The expression levels of *Col1a1* (**G**), *Col3a1* (**H**) and *ACTA2* (**I**) in control, STZ, STZ + GRo and control + GRo groups were determined by RT-qPCR. Data are presented as mean±SEM, n=8. **J-L**: Representative western blot images of TGF-β1 (**J**) and the phosphorylation levels of SMAD2 (**K**) /SMAD3 (**L**) in cardiac tissues from control, STZ, STZ + GRo and control + GRo groups **M**: Quantification of western blots of TGF-β1. n=6. **N**: Quantification of western blots of the phosphorylation levels of SMAD2. n=4. **O:** Quantification of western blots of he phosphorylation levels of SMAD3. n=4.

### GRo inhibits the aberrant cilia growth and downregulates cilia-specific genes in cardiac tissues of type 2 diabetic mice

To verify the roles of GRo in maintain ciliary homeostasis, the ciliary morphology and cilia-specific genes were assayed. The results of immunofluorescence staining revealed that cilia were abnormally elongated in the STZ group compared with the control group (***P<*0.0001, Fig. 4A–B**). The GRo treatment effectively inhibited the ciliary overgrowth in the STZ + GRo group relative to the STZ group (***P* =0.0044, Fig. 4A–B**). It has been reported that the TGF-β/SMAD signalling is closely associated with ciliary homeostasis and can be upregulated by PC1 encoded by *Pkd1*^[43]^. Therefore, cilia-specific genes (*Pkd1*, *Pkd2*, *KIF3A* and *IFT88*) in cardiac tissues were assayed. The results of western blot and RT-qPCR assays showed that the protein and mRNA levels of cilia-specific genes were significantly elevated in the STZ group compared with the control group (***P*<0.05, Fig.4C–J**). Compared with the STZ group, the GRo treatment in the STZ + GRo group markedly downregulated cilia-specific genes (***P*<0.05, Fig.4K–N**). It is suggested that GRo can maintain ciliary homeostasis by suppressing abnormal upregulation of cilia-specific genes in DiaCM.

**Fig. 4.**
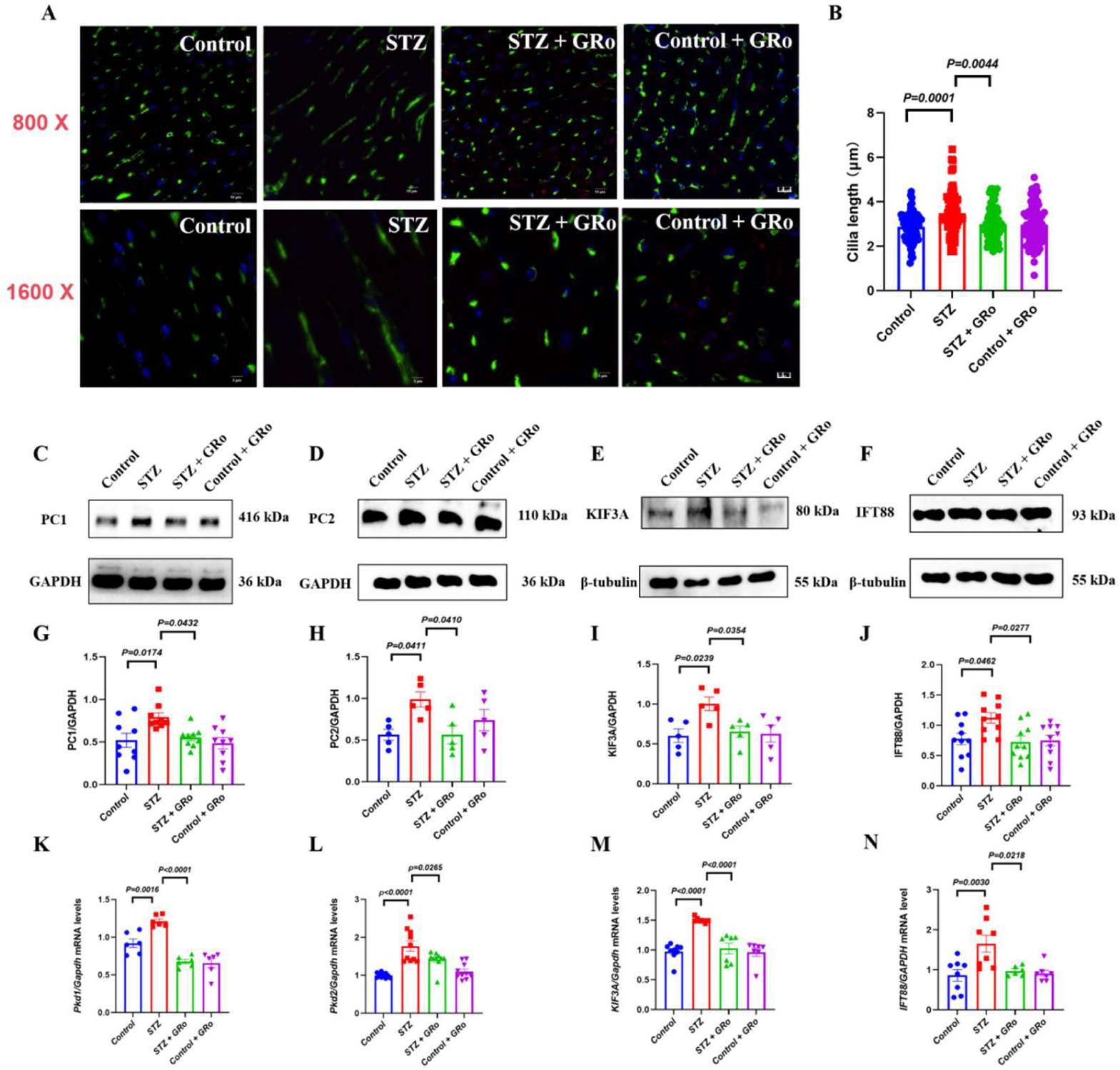
GRo inhibits the aberrant cilia growth and downregulates cilia-specific genes in cardiac tissues of type 2 diabetic mice. **A**: Representative fluorescence images of cilia in cardiac tissues from control, STZ, STZ + GRo and control + GRo groups of type 2 diabetic mice. Scale bars, 10 μm and 5 μm. **B**: Statistical graph of cilia length in cardiac tissues from control, STZ, STZ + GRo and control + GRo groups. Presented as mean±SEM, n=70. **C-F**: Representative western blot images of PC1 (**C**), PC2 (**D**), KIF3A (**E**), and IFT88 (**F**) in cardiac tissues from control, STZ, STZ + GRo and control + GRo groups. **G**: Quantification of western blots of PC1. n=9. **H**: Quantification of western blots of PC1. n=5. **I**: Quantification of western blots of PC1. n=5. **J**: Quantification of western blots of PC1. n=10. **K**: The expression levels of *Pkd1* in control, STZ, STZ + GRo and control + GRo groups was determined by RT-qPCR. Data are presented as mean±SEM. n=6. **L**: The expression levels of *Pkd2* in control, STZ, STZ + GRo and control + GRo groups was determined by RT-qPCR. Data are presented as mean±SEM. n =9. **M**: The expression levels of *KIF3A* in control, STZ, STZ + GRo and control + GRo groups was determined by RT-qPCR. Data are presented as mean±SEM. n=7. **N**: The expression levels of *ITF88* in control, STZ, STZ + GRo and control + GRo groups was determined by RT-qPCR. Data are presented as mean±SEM. n=8.

### GRo inhibits myocardial fibrosis induced by TGF-β1 in MCFs by downregulating the TGF-β/SMAD signalling

To verify the roles of GRo in myocardial fibrosis, we further examined the levels of fibrosis-related markers (COLI, COLIII and α-SMA) and the activation status of the TGF-β/SMAD signalling in MCFs. The results of western blot and RT-qPCR assays showed that the TGF-β1 stimulation significantly upregulated the protein and mRNA levels of the fibrosis-related markers in the TGF-β1 group compared with the control group (***P*<0.05, Fig.5A–I**). The GRo treatment in the TGF-β1 + GRo group markedly suppressed the overexpression of the fibrosis-related markers relative to the TGF-β1 group (***P* < 0.05, Fig. 5A–I**). Furtherly, the results of western blot assays showed that the phosphorylation of SMAD2/3 were significantly elevated in the TGF-β1 group compared with the control group (***P* < 0.05, Fig. 5J–M**), while the GRo administration in the TGF-β1+GRo group significantly inhibited the phosphorylation of SMAD2/3 compared with the TGF-β1 group (***P* < 0.05, Fig. 5J–M**). It is validated that GRo exerts anti-fibrotic effects by inhibiting the TGF-β/SMAD signalling *in vitro*.

**Fig. 5.**
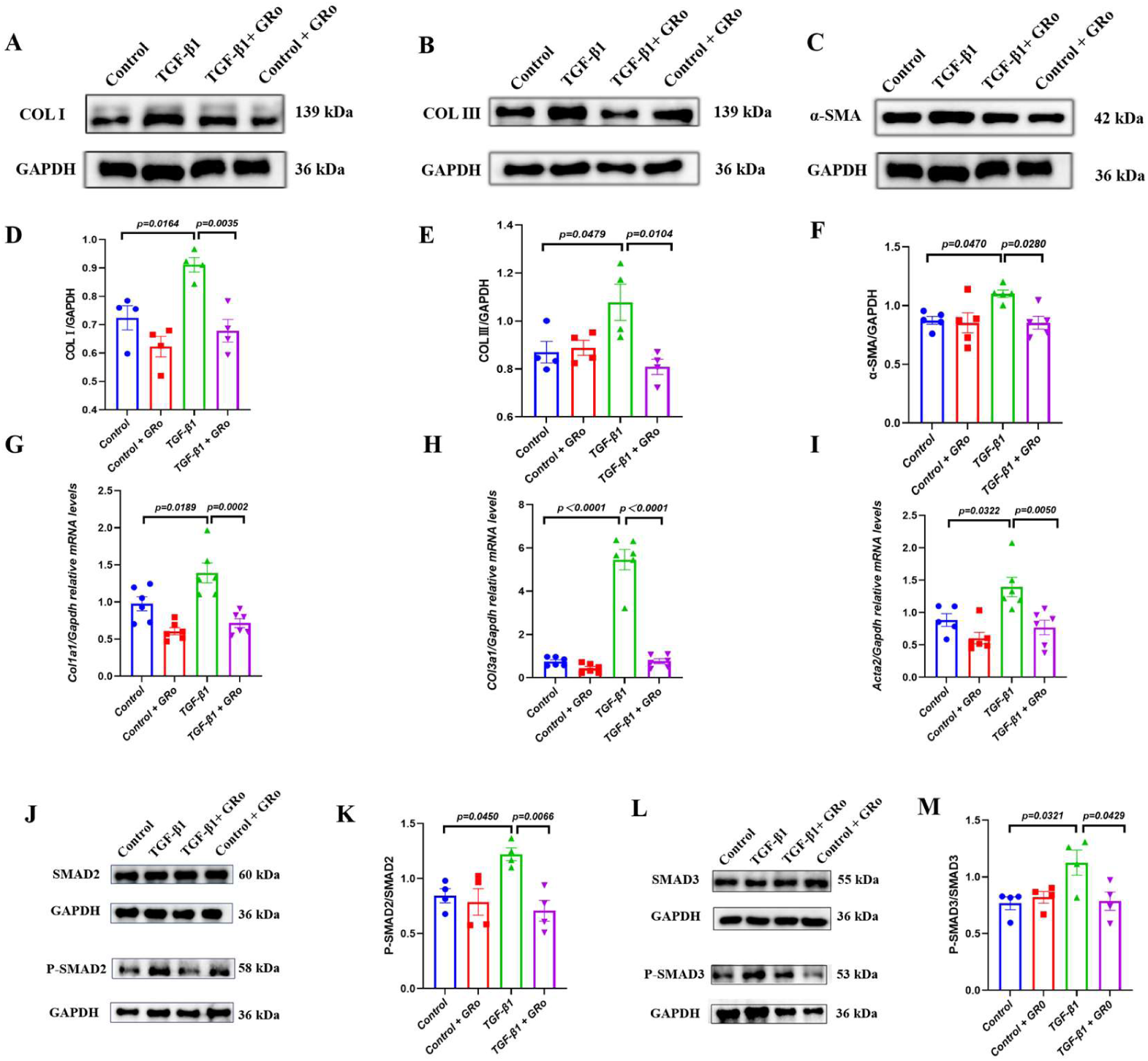
GRo inhibits myocardial fibrosis induced by TGF-β1 in MCFs by downregulating the TGF-β/SMAD signalling. **A-C**: Representative western blot images of COL I (**A**), COL III (**B**) and α-SMA (**C**) in MCFs from control, TGF-β1, TGF-β1 + GRo and control + GRo groups. **D**: Quantification of western blots of COL I. n=4. **E**: Quantification of western blots of COL III. n=4. **F**: Quantification of western blots of α-SMA. n=5. **G-I**: The expression levels of *Col1a1* (**G**), *Col3a1* (**H**) and *ACTA2* (**I**) in control, TGF-β1, TGF-β1 + GRo and control + GRo groups were determined by RT-qPCR. Data are presented as mean±SEM. n=6. **J-M**: Representative western blot images of the phosphorylation levels of SMAD2 (**J**) /SMAD3 (**L**) in MCFs from control, TGF-β1, TGF-β1 + GRo and control + GRo groups. Quantification of western blots of the phosphorylation levels of SMAD2 (**K**). n=4. Quantification of western blots of the phosphorylation levels of SMAD3 (**M**). n=4.

### GRo hinders the aberrant cilia growth and downregulates cilia-specific genes in MCFs

To verify the roles of GRo in maintain ciliary homeostasis, the ciliary morphology and cilia-specific genes were assayed. The results of immunofluorescence staining revealed that the TGF-β1 stimulation significantly promoted the cilia elongation in the TGF-β1 group compared with the control group (***P* < 0.05, Fig. 6A–B**). The GRo treatment effectively inhibited the ciliary overgrowth induced by TGF-β1 in the TGF-β1 + GRo group relative to the TGF-β1 group (***P* < 0.0001, Fig. 6A–B**). The results of western blot and RT-qPCR assays showed that the protein and mRNA levels of cilia-specific genes were significantly elevated in the TGF-β1 group compared with the control group (***P* < 0.05, Fig. 6C–N**). Compared with the TGF-β1 group, the GRo treatment in the TGF-β1 + GRo group markedly downregulated cilia-specific genes (***P* < 0.05, Fig. 6C–N**). In addition, it is indicated that the cilia-specific inhibitor HPI-4 significantly inhibited aberrant ciliary growth and myocardial fibrosis induced by TGF-β1, which suggests a close link between ciliary homeostasis and myocardial fibrosis as previous studies^[43]^ (***P* < 0.05, Supplementary Fig. 2A–E**). It is demonstrated that GRo suppresses aberrant ciliary growth by downregulating the overexpression of cilia-specific genes, which maintains ciliary homeostasis and ultimately attenuates myocardial fibrosis.

**Fig. 6.**
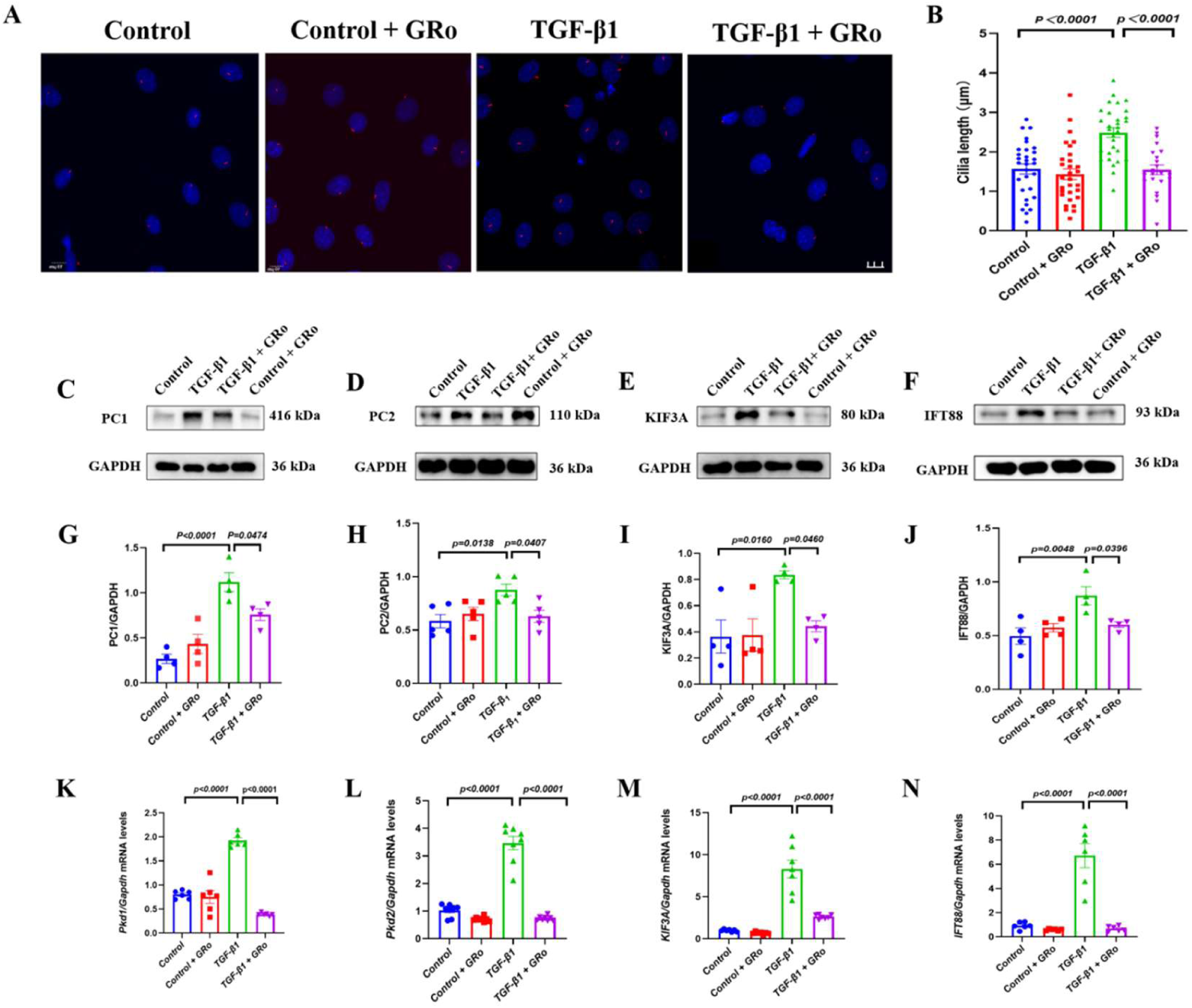
GRo inhibits the aberrant cilia growth and downregulates cilia-specific genes in MCFs. **A**: Representative fluorescence images of cilia in MCFs from control, TGF-β1, TGF-β1 + GRo and control + GRo groups. Scale bars, 10 μm **B**: Statistical graph of cilia length in MCFs from control, TGF-β1, TGF-β1 + GRo and control + GRo groups. Presented as mean±SEM. n =30. **C-F**: Representative western blot images of PC1 (**C**), PC2 (**D**), KIF3A (**E**), and IFT88 (**F**) in MCFs from control, TGF-β1, TGF-β1 + GRo and control + GRo groups. **G**: Quantification of western blots of PC1. n=4. **H**: Quantification of western blots of PC1. n=5. **I**: Quantification of western blots of PC1. n=4. **J**: Quantification of western blots of PC1. n=4. **K**: The expression levels of *Pkd1* in control, TGF-β1, TGF-β1 + GRo and control + GRo groups was determined by RT-qPCR. Data are presented as mean±SEM, n=6. **L**: The expression levels of *Pkd2* in control, TGF-β1, TGF-β1 + GRo and control + GRo groups was determined by RT-qPCR. Data are presented as mean±SEM. n =8. **M**: The expression levels of *KIF3A* in control, TGF-β1, TGF-β1 + GRo and control + GRo groups was determined by RT-qPCR. Data are presented as mean±SEM, n=7. **N**: The expression levels of *ITF88* in control, TGF-β1, TGF-β1 + GRo and control + GRo groups was determined by RT-qPCR. Data are presented as mean±SEM. n=6.

### The upregulation of *Pkd1* antagonizes the anti-fibrotic effect of GRo in MCFs

It is reported that the upregulation of *Pkd1* can lead to myocardial fibrosis by activating the TGF-β/SMAD signalling^[43]^. To further reveal the relationship between GRo and *Pkd1* in myocardial fibrosis, the *Pkd1*-targeted small activating RNA (sa*Pkd1*) was constructed, synthesized and transfected into MCFs. The results of screening experiments identified that 60 nM sa*Pkd1* significantly upregulated the expression level of *Pkd1* (***P* < 0.05, Fig. 7A–F**). It is revealed that the overexpression of *Pkd1* can inhibit the anti-fibrotic effect of GRo in TGF-β1-treated MCFs (***P* < 0.05, Fig. 7G–O**). The results of molecular docking analysis have shown that GRo can bind to PC1 with high affinity, which exhibits a binding energy of −8.282 kcal/mol (**Supplementary Fig. 3**). It is suggested that the PC1 may be the critical downstream target of GRo in preventing myocardial fibrosis.

**Fig. 7.**
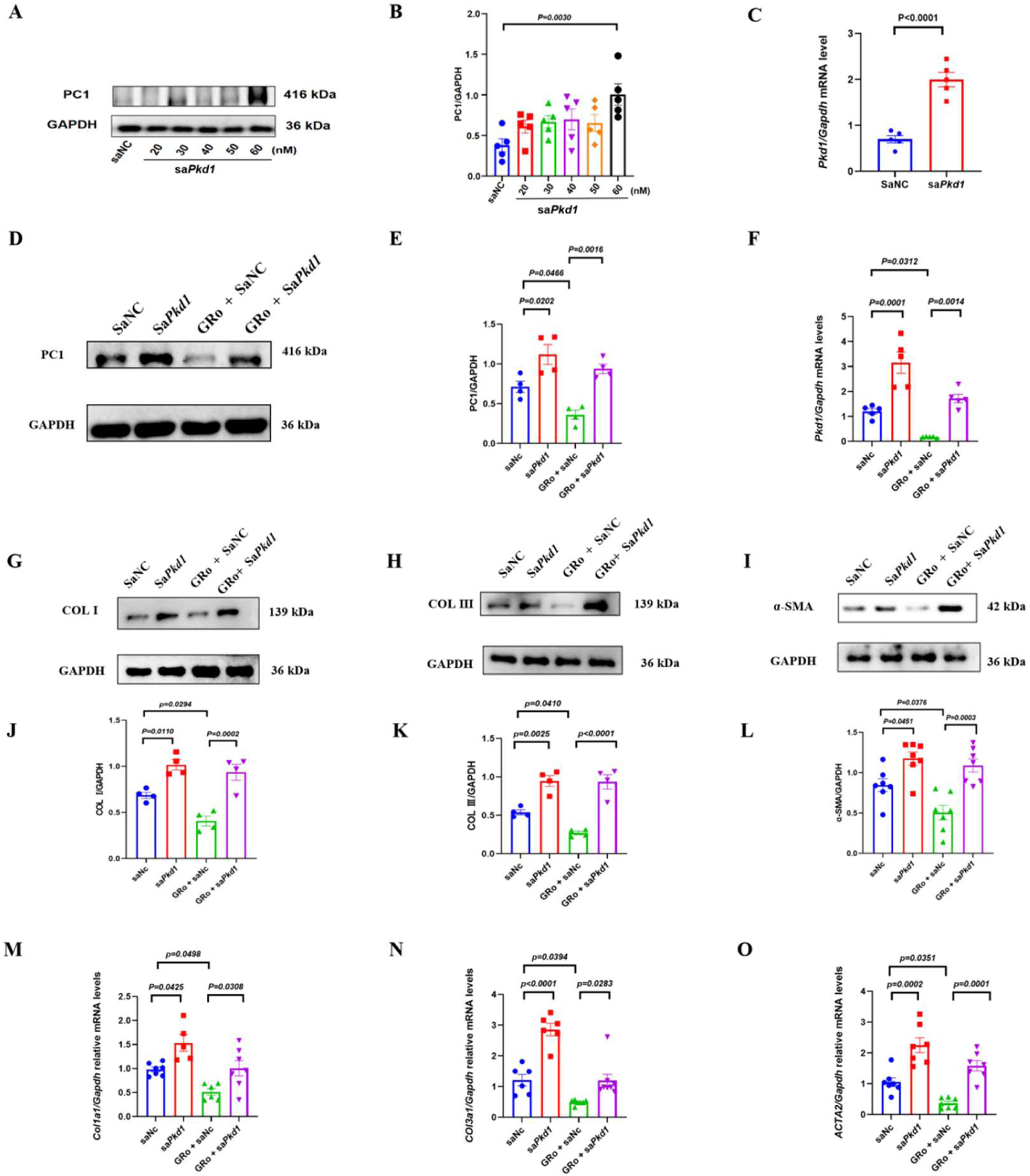
The upregulation of *Pkd1* antagonizes the anti-fibrotic effect of GRo in MCFs. **A**: Representative western blot images of overexpression of the *Pkd1* gene in MCFs at different concentrations. **B**: Quantification of western blots of overexpression of the *Pkd1* gene in MCFs at different concentrations. n=5. **C**: The expression levels of *Pkd1* in saNC and sa*Pkd1* groups were determined by RT-qPCR at a concentration of 60 nM. Data are presented as mean±SEM. n=5. **D**: Representative western blot images of PC1 in MCFs from saNC, sa*Pkd1*, GRo + saNC and GRo + sa*Pkd1* groups. **E**: Quantification of western blots of PC1 in MCFs from saNC, sa*Pkd1*, GRo + saNC and GRo + sa*Pkd1* groups. Data are presented as mean±SEM. n=4. **F**: The expression levels of *Pkd1* in saNC, sa*Pkd1*, GRo + saNC and GRo + sa*Pkd1* groups was determined by RT-qPCR. Data are presented as mean±SEM. n=5. **G-I**: Representative western blot images of COL I (**G**), COL III (**H**) and α-SMA (**I**) in MCFs from saNC, sa*Pkd1*, GRo + saNC and GRo + sa*Pkd1* groups. **G**: Quantification of western blots of PC1. n=4. **J**: Quantification of western blots of COL I. n=4. **K**: Quantification of western blots of COL III. n=4. **L**: Quantification of western blots of α-SMA. n=7. **M**: The expression levels of *Col1a1* in saNC, sa*Pkd1*, GRo + saNC and GRo + sa*Pkd1* groups was determined by RT-qPCR. Data are presented as mean±SEM. n=6. **N**: The expression levels of *Col3a1* in saNC, sa*Pkd1*, GRo + saNC and GRo + sa*Pkd1* groups was determined by RT-qPCR. Data are presented as mean±SEM. n=6. **O**: The expression levels of *ACTA2* in saNC, sa*Pkd1*, GRo + saNC and GRo + sa*Pkd1* groups was determined by RT-qPCR. Data are presented as mean±SEM. n=7.

### GRo alleviates myocardial hypertrophy in type 2 diabetic mice by enhancing antioxidant pathways

Besides of myocardial fibrosis, myocardial hypertrophy is also one of important pathological remodelling features of DiaCM^[22]^. The results of RT-qPCR assays showed that the fetal genes (*Nppa*, *Nppb* and *Myh7*) are reactivated in the STZ group compared with the control group (***P* < 0.0001, Fig. 8A–C**). The GRo treatment in the STZ + GRo group markedly suppressed the fetal genes relative to the STZ group (***P* < 0.0001, Fig. 8A–C**). It has been reported that the Nrf2-SLC7A11-GPX4 signalling is closely linked with the occurrence and development of myocardial hypertrophy^[52]^. The results of western blot and RT-qPCR assays showed that the expression levels of *Nrf2*, *SLC7A11* and *GPX4* were significantly downregulated while that of *P53* was significantly upregulated in the STZ group compared with the control group (***P* < 0.05, Fig. 8D–O**). The GRo treatment in the STZ + GRo group can enhance antioxidant pathways relative to the STZ group (***P* < 0.05, Fig. 8D–O**). It is indicated that GRo attenuates myocardial hypertrophy in type 2 diabetic mice by upregulating the Nrf2/SLC7A11/GPX4 antioxidant axis.

**Fig. 8.**
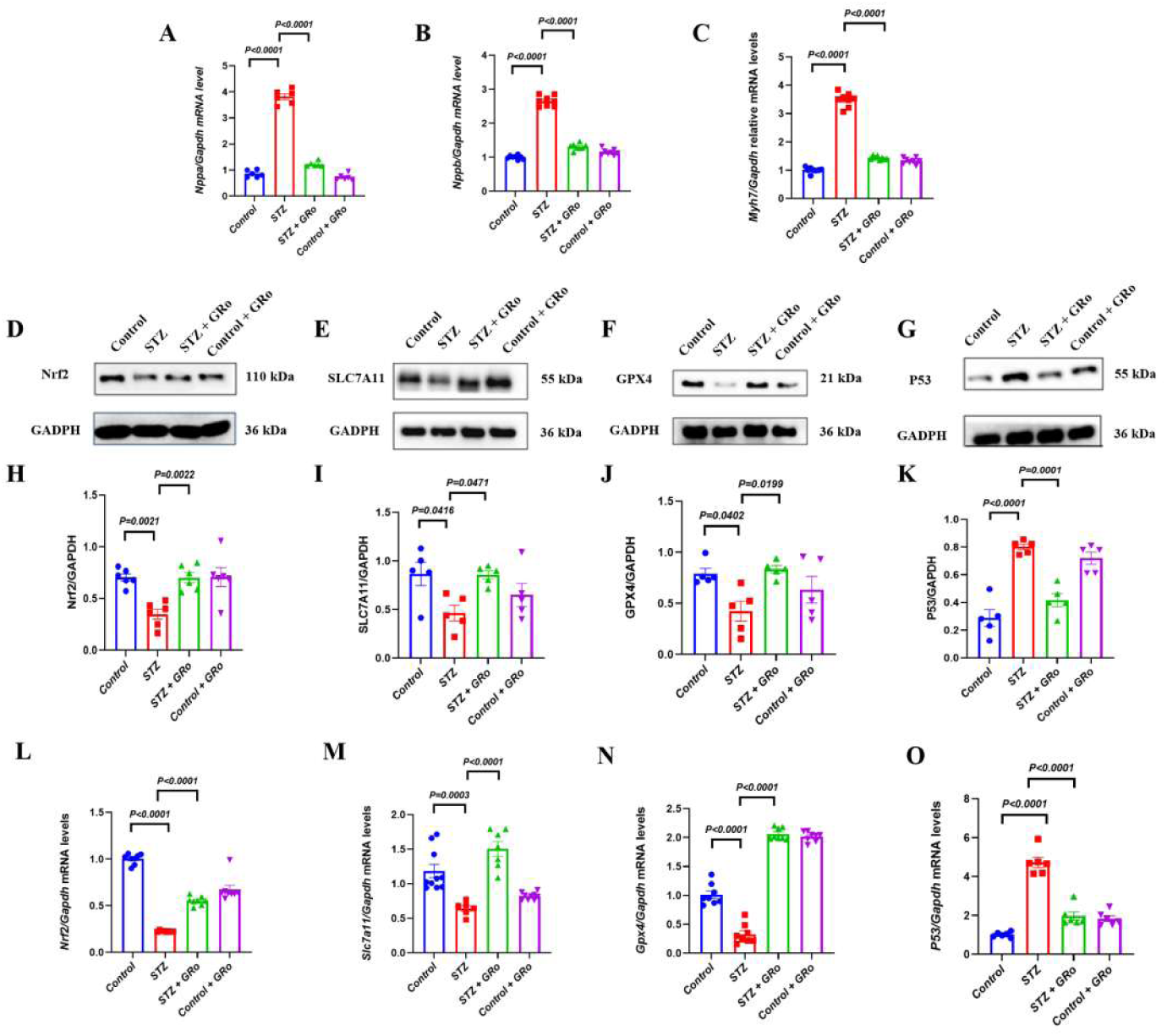
GRo alleviates myocardial hypertrophy in type 2 diabetic mice by enhancing antioxidant pathways. **A-C**: The expression levels of *Nppa* (**A**), *Nppb* (**B**) and *Myh7* (**C**) in control, STZ, STZ + GRo and control + GRo groups were determined by RT-qPCR. Data are presented as mean±SEM. n =7. **D-G**: Representative western blot images of Nrf2 (**D**), SLC7A11 (**E**), GPX4 (**F**), and P53 (**G**) in cardiac tissues from control, STZ, STZ + GRo and control + GRo groups. **H-K**: Quantification of western blots of Nrf2 (**H**), SLC7A11 (**I**), GPX4 (**J**) and P53 (**K**). n=5. **L**: The expression levels of *Nrf2* in control, STZ, STZ + GRo and control + GRo groups was determined by RT-qPCR. Data are presented as mean±SEM. n=8. **M**: The expression levels of *SLC7A11* in control, STZ, STZ + GRo and control + GRo groups was determined by RT-qPCR. Data are presented as mean±SEM. n=7. **N**: The expression levels of *GPX4* in control, STZ, STZ + GRo and control + GRo groups was determined by RT-qPCR. Data are presented as mean±SEM. n=8. **O**: The expression levels of *P53* in control, STZ, STZ + GRo and control + GRo groups was determined by RT-qPCR. Data are presented as mean±SEM. n=6.

### GRo inhibits cardiomyocyte hypertrophy by upregulating the Nrf2/SLC7A11/GPX4 antioxidant pathway

The results of immunofluorescence staining revealed that the palmitic acid (PA) stimulation significantly increased the cross-sectional area of cardiomyocytes and the results of RT-qPCR assays showed that the fetal genes were reactivated in the PA group compared with the control group (***P* < 0.05, Fig. 9A–E**). The GRo treatment in the PA + GRo group markedly suppressed the enlargement of cardiomyocytes and reactivation of fetal genes relative to the PA group (***P* < 0.001, Fig. 9A–E**). The results of western blot and RT-qPCR assays showed that the expression levels of *Nrf2*, *SLC7A11* and *GPX4* were significantly downregulated while that of *P53* was significantly upregulated in the PA group compared with the control group (***P* < 0.05, Fig. 9F–Q**). The GRo treatment in the PA + GRo group can enhance antioxidant pathways relative to the PA group (***P* < 0.05, Fig. 9F–Q**). In addition, it is revealed that the PA stimulation significantly upregulated the levels of reactive oxygen species (ROS) and malondialdehyde (MDA) while it downregulated the levels of glutathione (GSH) (***P* < 0.05, Fig. 9R–Z**). The GRo treatment in the PA + GRo group markedly suppressed oxidative stress response relative to the PA group (***P* < 0.01, Fig. 9R–Z**). The results *in vitro* are consistent with those *in vivo*, which further confirms that GRo can inhibit myocardial hypertrophy by enhancing antioxidant pathways.

**Fig. 9.**
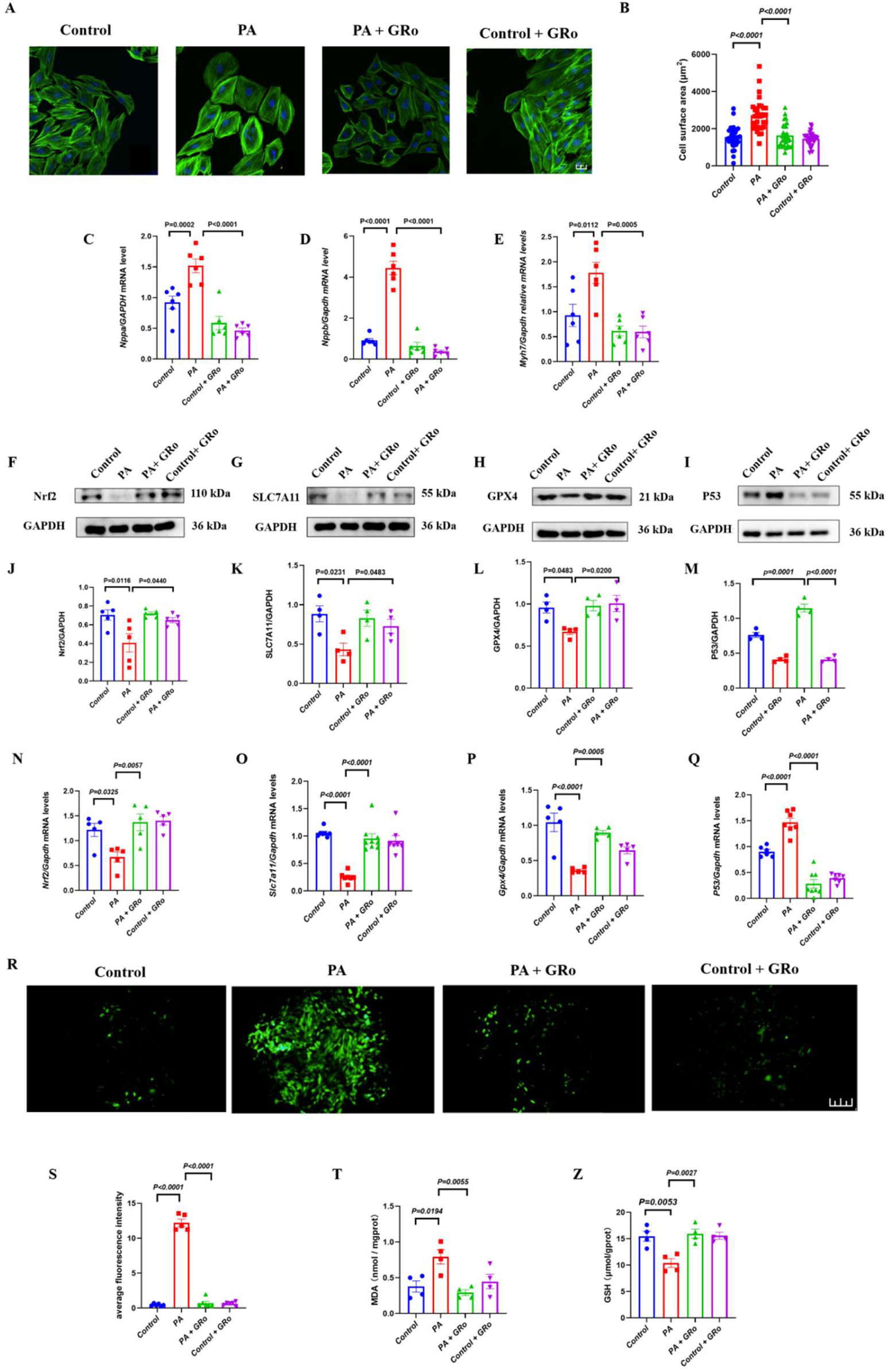
GRo inhibits cardiomyocyte hypertrophy by upregulating the Nrf2/SLC7A11/GPX4 antioxidant pathway. **A**: Representative fluorescence images of Cross-section of cardiomyocytes in H9c2 from control, PA, PA + GRo and control + GRo groups. Scale bars, 20 μm **B**: Statistical graph of Cardiomyocyte cross-sectional area in H9c2 from control, PA, PA + GRo and control + GRo groups. Presented as mean±SEM. n=30. **C-E**: The expression levels of *Nppa* (**C**)*, Nppb* (**D**) and *Myh7* (**E**) in control, PA, PA + GRo and control + GRo groups were determined by RT-qPCR. Data are presented as mean±SEM. n=6. **F-I**: Representative western blot images of Nrf2 (**F**), SLC7A11 (**G**), GPX4 (**H**), and P53 (**I**) in H9c2 from control, PA, PA + GRo and control + GRo groups. **J**: Quantification of western blots of Nrf2. n=5. **K-M**: Quantification of western blots of SLC7A11 (**K**), GPX4 (**L**) and P53 (**M**). n=4. **N**: The expression levels of *Nrf2* in control, PA, PA + GRo and control + GRo groups was determined by RT-qPCR. Data are presented as mean ± SEM. n=5. **O**: The expression levels of *SLC7A11* in control, PA, PA + GRo and control + GRo groups was determined by RT-qPCR. Data are presented as mean ± SEM. n = 9. **P**: The expression levels of *GPX4* in control, PA, PA + GRo and control + GRo groups was determined by RT-qPCR. Data are presented as mean ± SEM. n = 5. **Q**: The expression levels of *P53* in control, PA, PA + GRo and control + GRo groups was determined by RT-qPCR. Data are presented as mean ± SEM. n = 7. **R**: Representative fluorescence images of ROS of in H9c2 from control, PA, PA + GRo and control + GRo groups. Scale bars, 100 μm. **S**: Statistical graph of ROS in H9c2 from control, PA, PA + GRo and control + GRo groups. Presented as mean ± SEM. n = 5. **T-Z:** Relative levels of MDA and GSH in H9c2. Data are presented as mean ± SEM. n = 4.

## Discussion

Epidemiological data have indicated that the prevalence of diabetic cardiomyopathy (DiaCM) ranges from 30% to 60% among the approximately 463 million diabetic patients worldwide^[94-95]^. DiaCM is the leading cause of end-stage heart failure in diabetic patients and its clinical prevention and treatment is difficult^[96-97]^. It is defined as organic myocardial damage directly induced by diabetes independent of confounding factors such as coronary artery stenosis and hypertension, and its clinical course typically begins with early diastolic dysfunction, followed by myocardial interstitial fibrosis and ventricular hypertrophy, ultimately progressing to HFpEF^[98-101]^. The pathological process of DiaCM mainly involves pathological myocardial remodeling triggered by the sustained hyperglycemic microenvironment, which is consisted of myocardial fibrosis and hypertrophy^[102-107]^. The main classes of pharmacologic agents commonly used in clinical practice to modulate cardiac remodeling include angiotensin-converting enzyme inhibitors (ACEIs), angiotensin II receptor blockers (ARBs), beta-blockers and mineralocorticoid receptor antagonists^[108-110]^. They can partially reverse cardiac remodeling mainly through suppressing neurohormonal overactivation, reducing cardiac workload and exerting antifibrotic effects^[110-112]^. However, there are many limitations for them. They can exert side effects including hypotension, hyperkalemia, dry cough and bradycardia^[113-114]^. In addition, they cannot completely reverse myocardial fibrosis and hypertrophy, as well as have poor long-term medication adherence and substantial interindividual variability in treatment^[115-116]^. Therefore, unraveling the molecular mechanisms driving DiaCM and developing effective therapeutic agents have become pressing scientific imperatives.

Previous studies have reported that the oleanane-type ginsenoside Ro (GRo) exhibits anti-inflammatory, antioxidant, neuroprotective, vasoprotective, and anticancer effects in cardiovascular diseases while its roles in DiaCM remains unknown^[117-122]^. It is reveled that GRo can exert its cardioprotective effects and alleviate DiaCM through the dual-pathways in our studies (**Fig.10**). GRo plays a critical role in anti-myocardial fibrosis by maintaining ciliary homeostasis and downregulating PC1 *in vivo* and *in vitro* (**Fig.10**). In addition, it can enhance the antioxidant capacity by upregulating the Nrf2–SLC7A11–GPX4 axis, which inhibits myocardial hypertrophy *in vivo* and *in vitro* (**Fig.10**). Finally, it can ameliorate DiaCM. Nevertheless, there are still some aspects of this experiment that warrant further in-depth study. Firstly, the direct targets with which GRo interacts remain to be investigated. Secondly, the detailed mechanisms through which GRo regulates cilia-related genes and oxidative stress-related genes have yet to be elucidated. Finally, precise delivery methods of GRo into cardiac tissue remain to be developed.

**Fig. 10.**
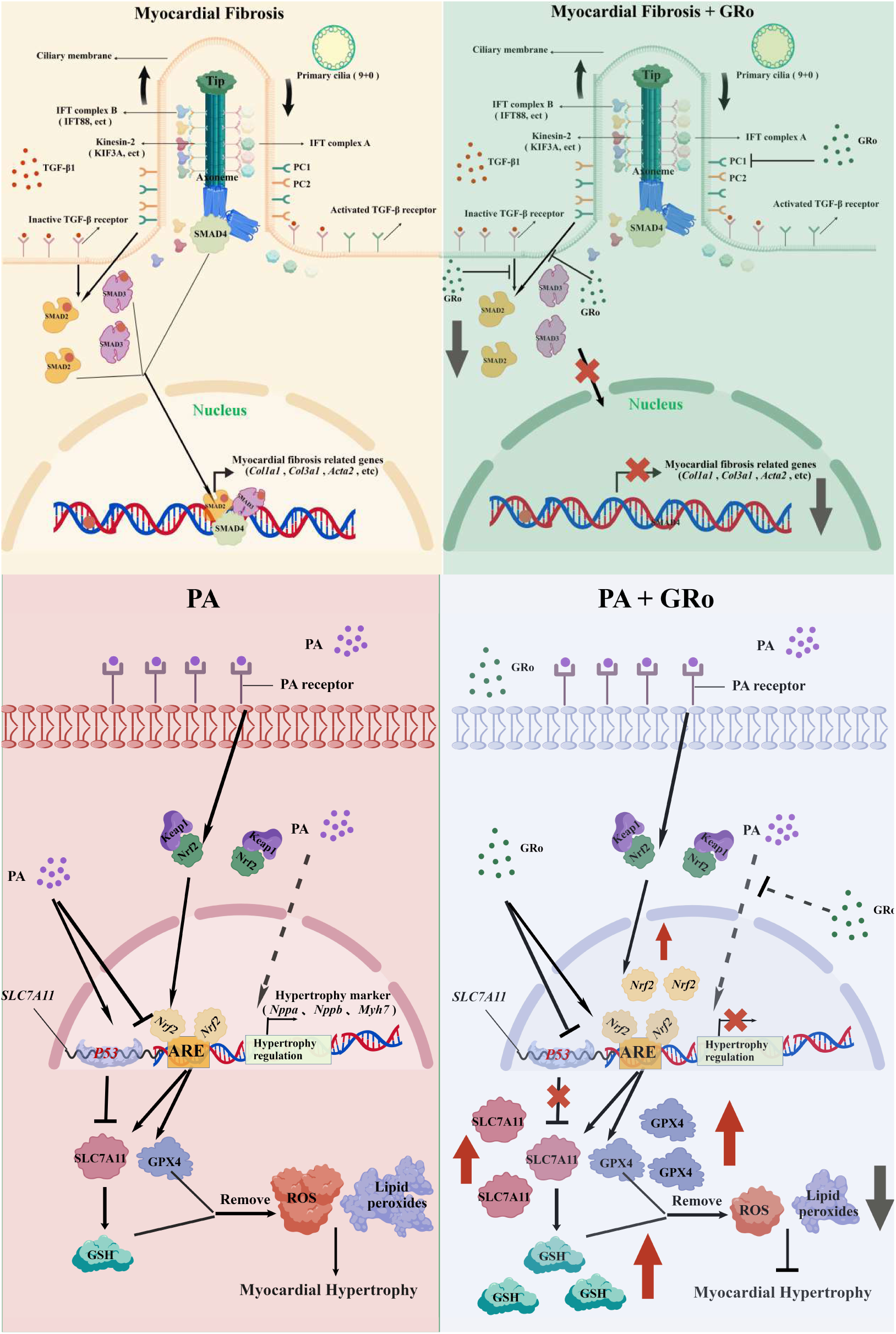
Mechanistic diagram of GRo in alleviating diabetic cardiomyopathy.

GRo, derived from a natural product, possesses a relatively low toxicity profile, reverses myocardial fibrosis and hypertrophy, improves long-term medication adherence, lowers blood glucose and exhibits minimal interindividual variability (**Fig. 1A**, **Fig. 3A-I**, **Fig. 5A-I**, **Fig. 8A-C**, **Fig. 9C-E**)^[123-124]^. Therefore, the combined administration of GRo and commonly used clinical Western drugs such as ACEIs and ARBs for the treatment of DiaCM may be a potential therapeutic strategy, which needs to be investigated in future. In addtion, it is reported that lipid nanoparticles (LNPs) have many advantages such as biocompatibility, facile surface functionalization and targeted delivery potential, which makes it become a safe and effective drug delivery vectors^[125-126]^. In future studies, LNPs may be employed to encapsulate and delieve GRo to myocardial tissue preciesly. Finally, dietary intervention during the preclinical or early stages of DiaCM carries unique value rooted in the concept of “preventive treatment of disease”^[127]^. Developing GRo into daily-use formulations such as functional foods, tea substitutes, solid beverages, or dietary supplements is expected to shift its therapeutic window to an earlier stage of disease, thereby providing a novel translational approach for the prevention and early intervention of diabetes or DiaCM in high-risk populations^[128-129]^.

## Availability of data and material

Not applicable.

## Code availability

Not applicable.

## Authors’ contributions

Z.Y. contributed to the methodology, carried out the experiments and wrote the manuscript. Y.G., X.G. and P.W. performed the animal experiments. B.G. and Y.S. conducted the cellular experiments. C.Z. and Y.T. analyzed the data. Z.R. designed the research, directed the study and wrote the manuscript.

## Ethics approval

Not applicable.

## Patient consent for publication

Not applicable.

## Consent to participate

All the authors have confirmed their participation.

## Consent for publication

All the authors agree to the publication of the current version of the manuscript.

## Acknowledgments

We are grateful to all the participants of this work.

## Supplementary Diagram

**Supplementary Fig. 1.**
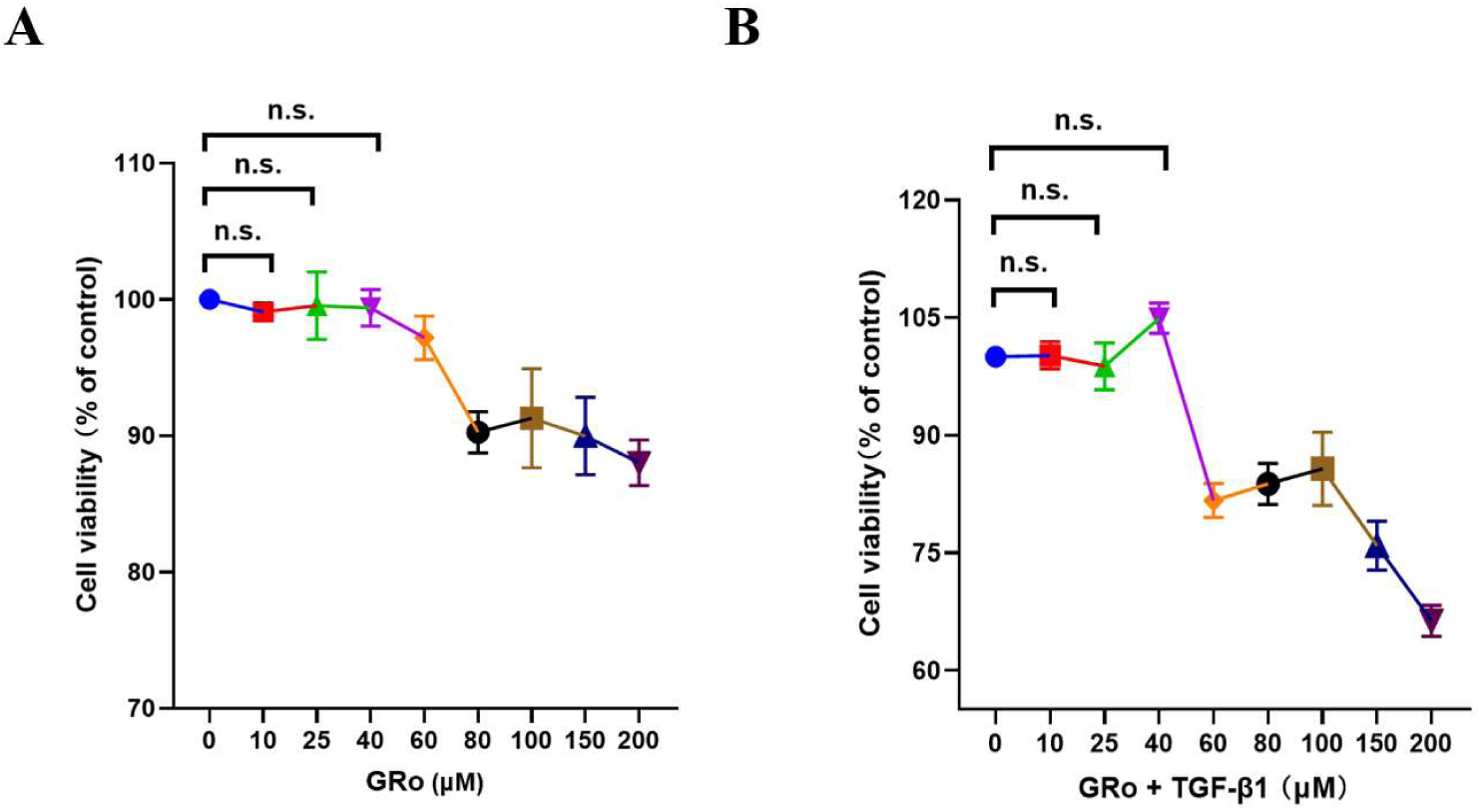
The effect of GRo and combined effect of GRo with TGF-β1 in MCFs. **A**: Effect of GRo with different concentrations in MCFs. Data are presented as mean ± SEM. n = 6. **B:** Effect of GRo combined with TGF-β1 in MCFs. Data are presented as mean ± SEM. n = 6.

**Supplementary Fig. 2.**
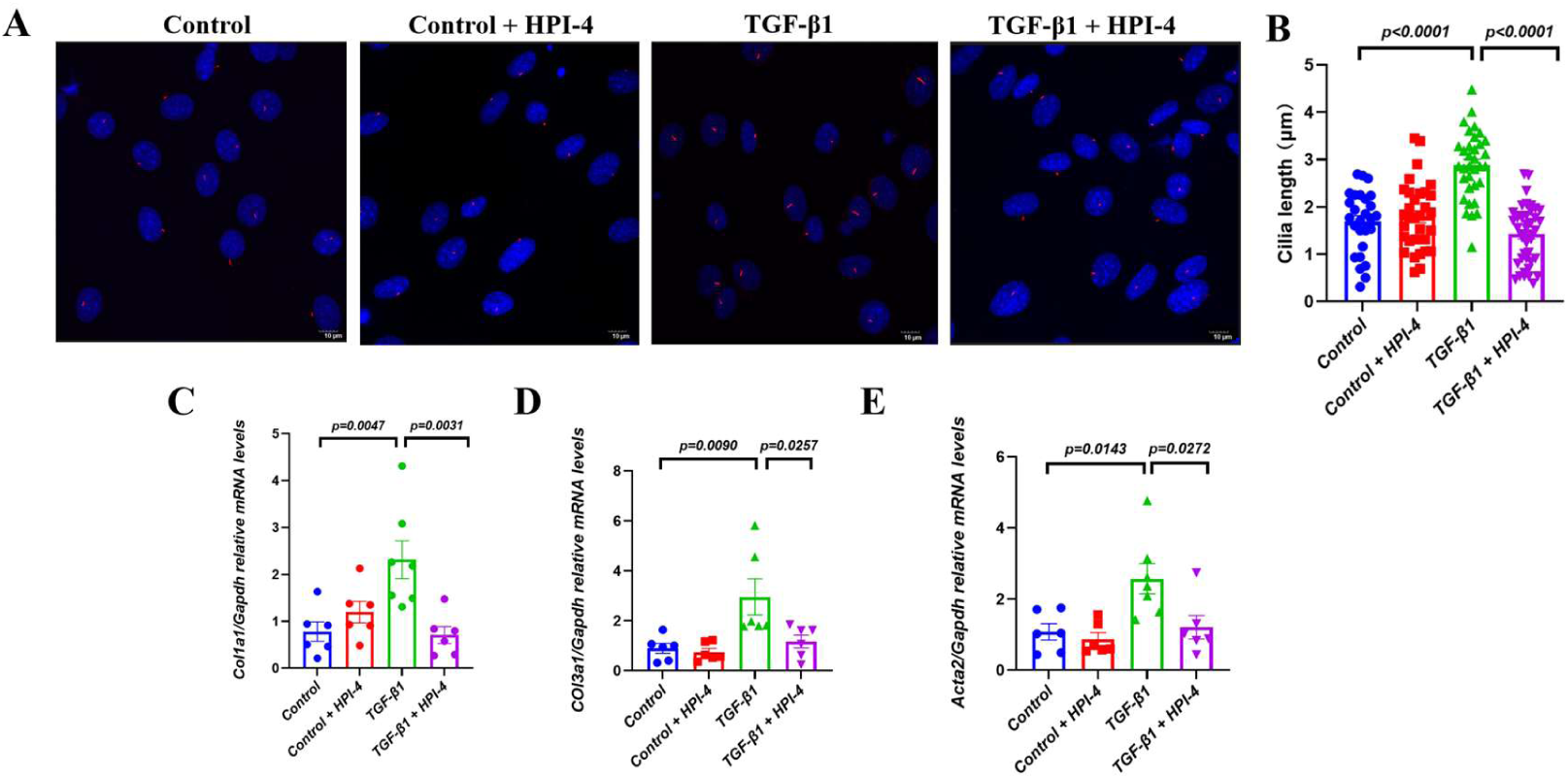
HPI-4 attenuates fibrosis in MCFs by inhibiting aberrant cilia growth. **A**: Representative fluorescence images of cilia in MCFs from control, TGF-β1, TGF-β1 + HPI-4 and control + HPI-4 groups. Scale bars, 10 μm **B**: Statistical graph of cilia length in MCFs from control, TGF-β1, TGF-β1 + HPI-4 and control + HPI-4 groups. Presented as mean ± SEM. n = 30. **C-E**: The expression levels of *Col1a1* (**C**), *Col3a1* (**D**) and *ACTA2* (**E**) in control, TGF-β1, TGF-β1 + HPI-4 and control + HPI-4 groups were determined by RT-qPCR. Data are presented as mean ± SEM. n = 6.

**Supplementary Fig. 3.**
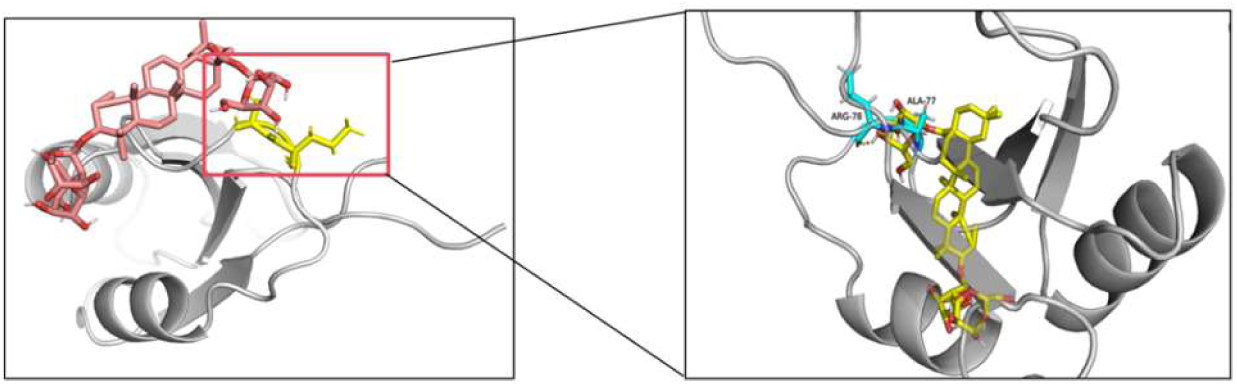
Molecular docking analysis of GRo and PC1. Left panel shows the overall docking conformation, and right panel shows a detailed view of the binding interface. The binding affinity between GRo and PC1 was −8.282 kcal/mol. The small molecule forms hydrophobic interactions with ALA77 and establishes two distinct hydrogen bonds with ARG78 and ALA77, respectively.

**Supplementary Fig. 4.**
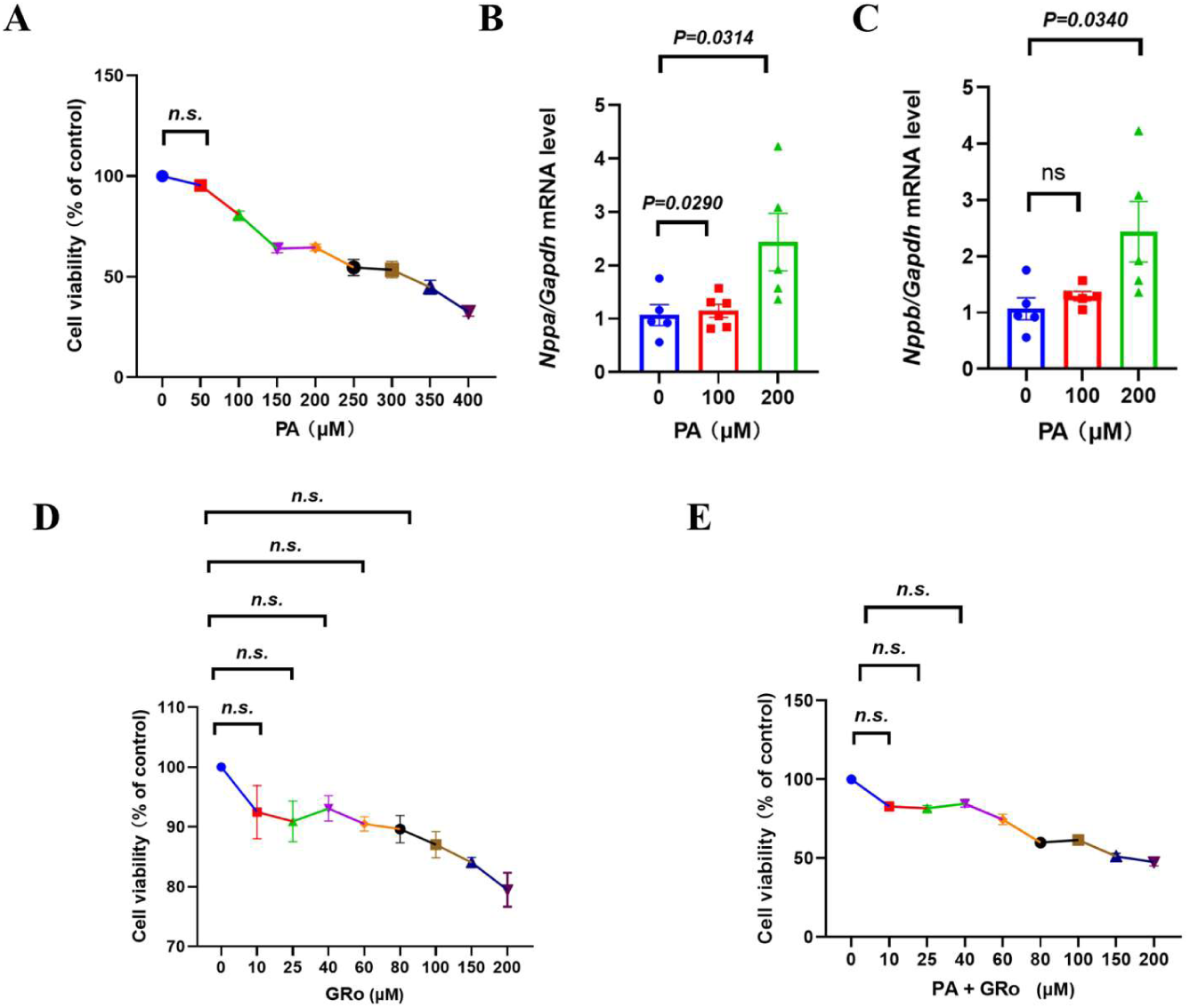
The effect of PA and combined effect of GRo with PA in cardiomyocyte. **A**: Effect of PA with different concentrations in the viability of cardiomyocyte. Data are presented as mean ± SEM. n = 6. **B–C**: The expression levels of *Nppa* (**B**) and *Nppb* (**C**) at 100 and 200 μM PA were determined by RT-qPCR. Data are presented as mean ± SEM. n = 5. **D**: Effect of GRo with different concentrations in the viability of cardiomyocyte. Data are presented as mean ± SEM. n = 6. **E**: Effect of GRo combined with PA in the viability of cardiomyocyte. Data are presented as mean ± SEM. n = 6.

**Supplementary Table 1.**
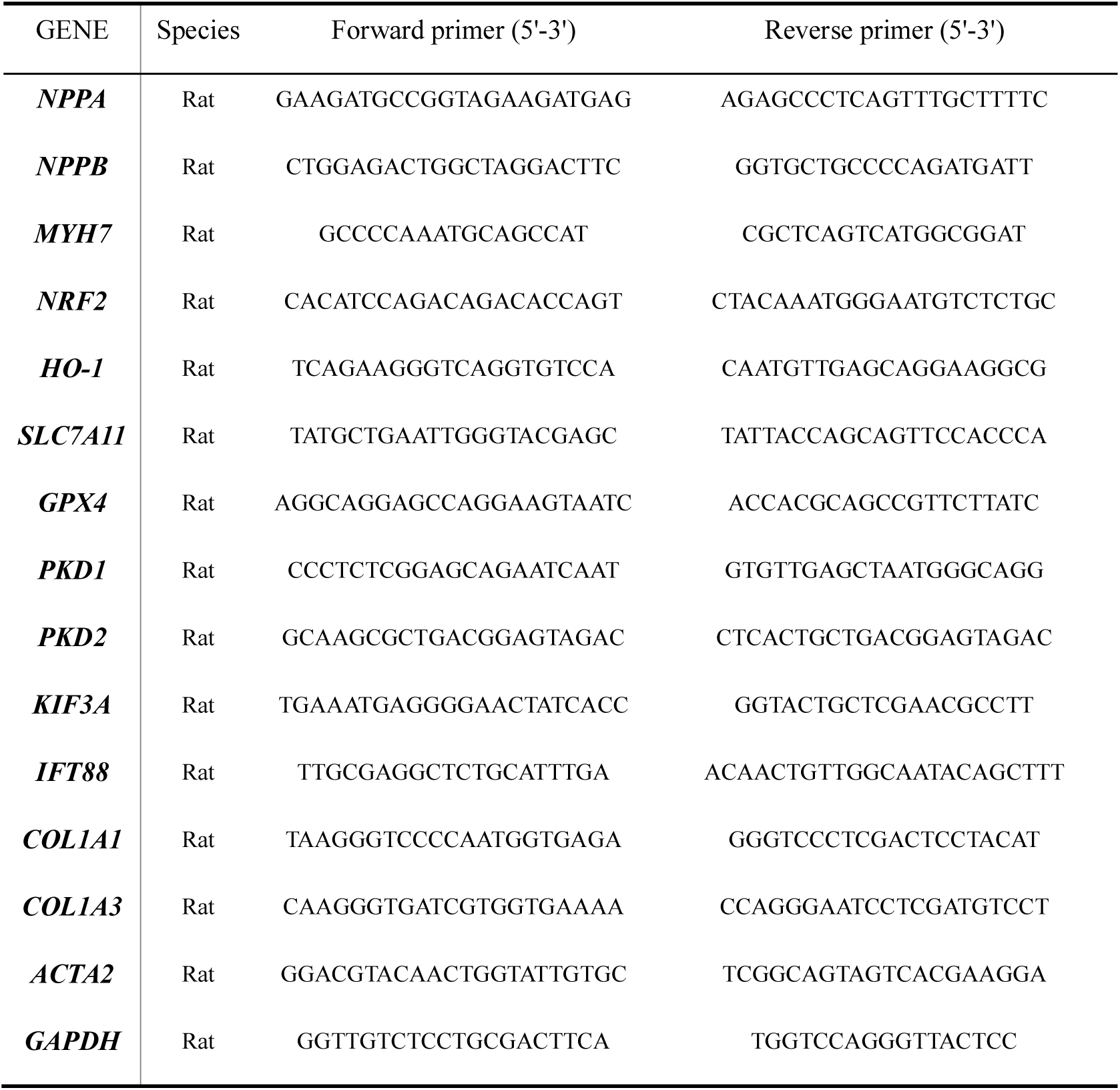
The relevant primer sequences.

**Supplementary Table 2.**
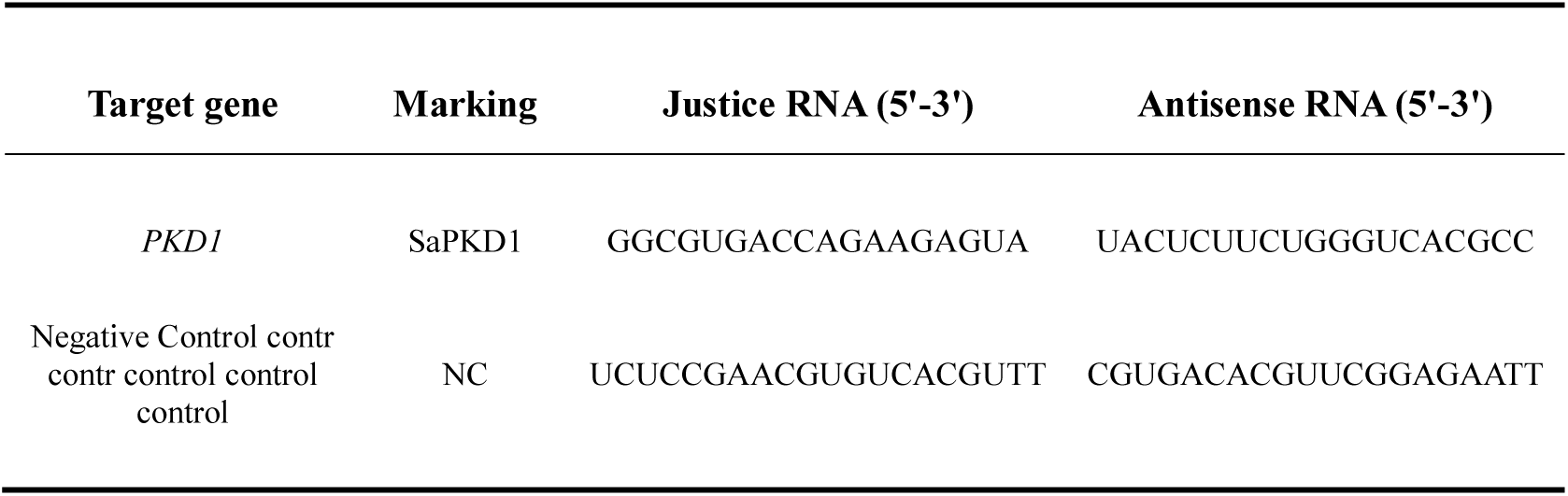
The saRNA-related sequences in the study.

